# A Yeast Two-Hybrid Protein Domain Screening Approach for Ebola Virus-Human Protein Interactions Identifies PABPC1 as a Host Factor Required for Replication

**DOI:** 10.64898/2026.03.05.709814

**Authors:** James P. Connell, Venkatesh Sivanandam, Sarah Hulsey Stubbs, Callie J. Donahue, Veronica J. Heintz, Bruno La Rosa, Olena Shtanko, Robert A. Davey, Douglas J. LaCount

## Abstract

Ebola virus (EBOV) causes severe hemorrhagic fever and remains a global health threat despite advances in vaccines and antibody-based therapeutics. Viral replication and immune evasion depend on interactions between EBOV proteins and host factors, yet the full spectrum of these interactions is unknown. To systematically identify EBOV-human protein-protein interactions, we performed more than 200 yeast two-hybrid (Y2H) screens using full-length EBOV proteins and domain fragments as baits and cDNA libraries derived from human liver tissue, macrophages, and interferon-stimulated macrophages. This approach revealed 521 unique interactions supported by confidence scoring based on retest assays and library screen data. The interactome includes proteins involved in RNA metabolism, transcriptional regulation, and post-translational modification, with enrichment for RNA-binding proteins among partners of NP, VP30, and L. By employing multiple constructs for each EBOV gene, interaction interfaces could be localized to specific viral domains. This approach identified a likely MYND domain binding site in the intrinsically ordered region of NP. Similarly, overlapping cDNA fragments from Y2H screens revealed potential binding sites within human proteins, including putative VP30 binding sites on NUFIP2 and PABPC1 that lacked canonical PPPPxY motifs. PABPC1 interacted with VP30 in uninfected human cells and accumulated in inclusion bodies in infected cells that resembled respiratory syncytial virus inclusion body associated granules (IBAGs). Inhibiting PABPC1 expression reduced EBOV replication early in infection. These findings expand the EBOV-host interaction network, identify candidate regulators of viral RNA synthesis, and provide a resource for mechanistic studies and antiviral target discovery.

**Importance:** Ebola virus (EBOV) remains a major global health threat due to its high mortality rate and limited treatment options. Viral replication and immune evasion depend on interactions between EBOV proteins and host factors, yet these interactions are incompletely understood. This study uses a large-scale yeast two-hybrid approach to systematically map EBOV-human protein-protein interactions, revealing hundreds of previously unreported partners and identifying cellular pathways that may be exploited by the virus. By integrating domain mapping, confidence scoring, and functional validation, we provide a resource that advances understanding of EBOV biology and may be used to discover candidates for antiviral development.

## Introduction

Ebola virus (EBOV), a member of the *Filoviridae* family, is a non-segmented, negative-sense RNA virus that causes severe hemorrhagic fever in humans and nonhuman primates. The largest EBOV outbreak to date caused more than 11,000 deaths in West Africa from 2013-2016, but smaller outbreaks continue to occur sporadically, with the most recent in Democratic Republic of the Congo during the fall of 2025 (1). The continued emergence of EBOV highlights the urgent need for a better understanding of the EBOV replication and improved therapeutic strategies. Although vaccines and monoclonal antibody treatments have been developed, these interventions are limited by cost, accessibility, and timing of administration (1). A deeper understanding of EBOV biology, including the molecular interactions between viral and host proteins, will help to identify new targets for antiviral development.

The EBOV genome encodes seven major proteins that are primarily responsible for viral genome replication, transcription of viral mRNAs, and assembly and release of infectious viral particles. Nucleoprotein (NP), polymerase cofactor VP35, and the RNA-dependent RNA polymerase protein L form the protein complex that orchestrates replication of the viral RNA genome within cytoplasmic inclusion bodies (2–4). Adding VP30 to the complex enables transcription of viral genes (4–6). NP, together with VP24 and VP35, form the filamentous nucleocapsid structures that encapsulate viral RNA (7–13). The matrix protein, VP40, surrounds nucleocapsids and mediates viral particle assembly and budding from the host cell (14–16). As particles bud, they acquire a lipid membrane in which the envelope glycoprotein, GP, is embedded. GP mediates entry through interactions with host cell receptors and fusion with cellular membranes (17). Although these processes are predominantly driven by viral factors, host proteins play critical roles. For example, VP30 is phosphorylated by the cellular protein kinase SRPK1, which reduces binding to viral RNA and suppresses transcription (18). NP counters this effect by recruiting host phosphatase complexes to dephosphorylate VP30, which increases VP30 binding to RNA and stimulates transcription (19, 20). RBBP6 adds an additional level of regulation by competing with NP for binding to VP30 (21, 22). In addition, EBOV proteins, in particular VP24 and VP35, interact with multiple pathways to thwart the host immune response (23). Yet despite significant progress over the past decade, much remains to be discovered about the interactions between EBOV and host proteins.

Systematic mapping of virus-host PPIs is a powerful strategy to uncover cellular pathways exploited by EBOV. Previous studies have used affinity purification coupled with mass spectrometry (AP-MS) and proximity dependent biotinylation to identify host interactors of EBOV and cellular proteins (21, 24–31). While these approaches provide valuable insights, they often capture multiprotein complexes rather than direct binary interactions, making it difficult to distinguish primary contacts from indirect associations. Furthermore, there is little overlap in the interactions identified between the studies, which is a general limitation of the field and is true for any examination of physical or functional virus-host interactions. There is currently no single approach that can identify all protein-protein interactions (PPIs), and differences in experimental conditions, instrumentation, constructs, and cell lines employed further exacerbate the issue. Only by performing multiple studies using different approaches can the complete set of interactions be identified for any system.

Here we implemented the yeast two-hybrid (Y2H) assay to screen for EBOV-human protein interactions. In this system, EBOV proteins were expressed in yeast as fusions to the Gal DNA binding domain and screened against cDNA libraries of human gene fragments fused to the Gal4 transcription activation domain. Binding of an EBOV protein to a human protein generates a functional Gal4 transcription factor, which stimulates expression of Y2H reporter genes that enable growth on selective medium. The Y2H assay can be employed in high throughput screens of complex libraries encoding host proteins with no need for specialized containment facilities or complex purification steps. It usually detects direct interactions between two proteins and can reveal weak or transient interactions that may escape detection by biochemical methods. Thus, genome-wide Y2H screens complement co-affinity-purification and proximity dependent biotinylation for the identification of virus-host interactions.

In this study, we performed more than 200 yeast two hybrid screens of full-length EBOV proteins and protein domains against cDNA libraries from human liver tissue, macrophages and interferon-stimulated macrophages. We identified more than 500 putative EBOV-human protein interactions with confidence scores for each interaction. The resulting dataset offers new insights into the molecular interface between EBOV and its human host and establishes a resource for future investigations into virus-host interplay.

## Materials and methods

### Preparation of activation domain libraries

Monocytes were isolated from the blood of healthy donors collected according to University of Texas Health-approved IRB protocol 12-05-5536 and differentiated into macrophages as previously described (32). Following differentiation, recombinant interferon-β was added to one set of macrophages for 18 hours to induce expression of interferon stimulated genes (ISGs). Total RNA was isolated from untreated and interferon-β-treated macrophages using TRIzol LS Reagent (Invitrogen, #10296028) according to the manufacturer’s protocol. Human Reference Total RNA, a mixture of total RNA from five human tissues, was purchased from TaKaRa Clontech (#636690). RNA quality was assessed on an Agilent Bioanalyzer 2100. All samples had an RNA Integrity Number (RIN) of at least 7.9.

Unidirectional cDNA was prepared from 1 μg of total RNA using the NEBNext® Ultra™ Directional RNA Library Prep Kit for Illumina following the “Protocol for use with NEBNext Poly(A) mRNA Magnetic Isolation Module (NEB #E7490)” through step 1.7 (Manual Version 5.0 5/15) with the following changes: 1) poly(A)+ mRNA was eluted for 5 minutes at 65 °C to prevent fragmentation; 2) after synthesizing second strand cDNA, the cDNA was purified using a Zymo Research DNA Clean & Concentrator-5 column (#D4013); 3) the adaptor primer (/5Phos/GATCGGAAGAGCTTGTTCTACCGAGGGACCC/ideoxyU/ ACTACTGCCTAAC GAACTCCCGCTCTTCCGATC*T; * indicates a phosphorothioate bond; synthesized and PAGE purified by IDT) was a modification of the NEBNext Adaptor for Illumina (E7352A). Following adaptor ligation, the DNA was purified using a Zymo Research DNA Clean & Concentrator-5 column and amplified with KAPA HiFi HotStart PCR polymerase (#KK2502) in a 50 μl reaction containing 3 μl of the USER enzyme (Uracil-Specific Excision Reagent, a mixture of uracil DNA glycosylase and endonuclease VIII) from the Library Prep Kit and primers 5’-GGGTCCCTCGGTAGAACAAGCTCT-3’ and 5’-ACTACTGCCTAACGAACTCCCGCTC-3’. The reaction was incubated 15 min at 37 °C to enable the USER enzyme to create single nucleotide gaps in the second strand cDNA and in the adaptor oligo, followed by 3 minutes at 98 °C for and 15 cycles of 98 C for 20 seconds, 60 C for 15 seconds, and 72 C for 30 seconds with a 1-minute extension at 72 °C at the end. The PCR products were purified using a DNA Clean & Concentrator-5 column and size fractionated on a Sephacryl S500 (GE Healthcare, #17-0613-10) column prepared in a disposable 2-ml plastic pipette (33). cDNA libraries were prepared with PCR products from fractions 5 plus 6 (700 - ∼2000 bp) and fraction 7 (500 – 1500 bp). Libraries from fractions 5 plus 6 were dominated by a common false positive and were not used for yeast two-hybrid library screens.

To generate yeast two-hybrid libraries, cDNAs were PCR amplified with KAPA HiFi HotStart PCR polymerase using primers 5’-CTAGTTCTGGTAGCTCTAAAGGGTCCCTCGGTAGAACAAGCTCT-3’ and 5’-TGTGACCACTTCCCGAACTACTGCCTAACGAACTCCCGCTC-3’ to add the 5’ extensions needed for recombination into the activation domain plasmid pOAD102 (33, 34). Twenty cycles were performed using the conditions described above. PCR products were co-transformed with linear pOAD102 into yeast strain BK100 (MAT*a* ura3-52 ade2-101 trp1-901 leu2-3,112 his3-200 gal4Δ gal80Δ GAL2-ADE2 LYS2::GAL1-HIS3 met2::GAL7-lacZ). Yeast were selected on synthetic drop-out (SD) medium lacking leucine and uracil (SD-LU) supplemented with 1.5X adenine (1X is 0.02 g/L). Colonies were collected in YEP (Yeast extract-peptone medium) to which 5% (v/v) DMSO was added. Aliquots of the library were stored at -80 °C until use. Titers were determined by thawing an aliquot of the frozen cultures and plating ten-fold dilutions on SD-LU supplemented with 1.5X adenine.

### Bait generation

EBOV Y2H DNA binding domain constructs (referred to as BD or bait) were generated in plasmid pOBD7A (35). Both full length genes and gene fragments corresponding to known domains or domains predicted based on sequence homology to related viral proteins of known structure were cloned. Two plasmids were generated for each viral gene or gene fragment, one in which the viral gene was inserted at the 5’ end of the GAL4 binding domain (designated pOBD7N) and one in which the viral gene was inserted at the 3’ end (designated pOBD7C). To generate pOBD7N and pOBD7C constructs, pOBD7A was digested with *Stu*I plus *Spe*I or *Eco*RI plus *Pst*I, respectively. EBOV genes and gene fragments were amplified from plasmids encoding full length genes with primers that encode 5’ sequences homologous to the flanking regions at sites of linearization for pOBD7N or pOBD7C (Table S1). PCR products were inserted into pOBD7A by homologous recombination in R2HMet (MATα ura3-52 ade2-101 trp1-901 leu2-3,112 his3-200 met2Δ::hisG gal4Δ gal80) (34). Colonies with the correct size insert were identified by PCR and sequenced.

### Self-activation of binding domain clones

R2HMet strains containing BD-viral gene plasmids (BD strains) were mated to a BK100 strain containing pOAD102-GFP, in which GFP was inserted in frame with the GAL4 AD and the *S. pombe URA4* gene. Diploid yeast were selected on SD-LU plates. A single colony was picked, diluted in sterile water, spotted on Y2H selection medium (SD lacking TRP, LEU, URA, HIS, and adenine (SD-TLUHA) supplemented with 1 mM, 2.5 mM, 5 mM, or 10 mM 3-amino-1,2,4-triazole (3-AT)), and incubated for 5-7 days at 30°C. The lowest concentration of 3-AT that suppressed yeast growth was used in subsequent yeast two-hybrid screens. Bait constructs that yielded growth at 10 mM 3-AT were not screened.

### Y2H library screens

Y2H screens were performed as described with slight modifications (33, 36). BD strains were grown overnight in liquid SD-T supplemented with 1.5X adenine at 30° C with rotation. The next morning, cultures were diluted to an OD_600_ of 0.4 in yeast extract peptone dextrose (YPD) medium supplemented with 1.5X adenine (YPD+A) and incubated ∼ 3 h at 30° C with rotation until the cultures reached an OD600 between 1.0 and 1.5. BD strains were then mated to the AD library in BK100 by adding 1 × 10^7^ cfu of each strain to a sterile culture tube, collecting the cells by centrifugation, resuspending in 3 ml YPD pH 3.5 and incubating for 90 min at 30° C with rotation. The yeast were then collected by centrifugation, resuspended in YPD pH 4.5 and subjected to centrifugation once more to pellet the yeast at the bottom of the tube. Culture supernatants were not removed. The cultures were incubated 4 h at room temperature without rotation. After the incubation, the cultures were resuspended by gentle vortexing and an aliquot removed to estimate the number of diploids formed by growth on solid SD-TLU+A. The remaining yeast were collected by centrifugation, washed once with water, resuspended in water, plated on solid SD-TLUHA plus 3-AT at the concentration previously determined to suppress background growth and incubated at 30° C. Y2H selections were performed on media lacking histidine and adenine because our initial test screens found IDO1 to be a common contaminant in most screens when the selections were performed on medium lacking histidine only.

However, the number of IDO1-containing colonies was suppressed when the Y2H screens were performed on medium lacking both histidine and adenine. Single colonies were picked between days 4-10 after mating and used to inoculate 100 μl of YPDA in a well of a sterile 96 well plate. Cultures were incubated overnight at 30° C without rotation or agitation. Yeast were resuspended by gentle vortexing and 10 μl were removed for PCR; 5 μl of DMSO was added to the remaining yeast and the cultures were frozen at -80° C.

To identify the human gene inserted into the AD plasmid, 40 μl of 2.5 mg/mL zymolyase (Amsbio LCC, Cambridge, MA, USA) dissolved in 100 mM sorbitol, 10 mM Tris, pH 7.5 was added to the 10 μl of yeast culture and the mixture incubated 10 min at 37° C and then 5 min at 95° C. AD inserts were amplified using forward primer 5’-CGACGACGAGGATACGCCACCGAAC-3’ and either reverse primer 5’-CCTCTGCCCTCGCCGAATAGCTCTG-3’ (for macrophage, interferon-treated macrophage, and universal libraries) or 5’-GAGCTTCGCAGCAACCGGACTAGGA-3’ (for liver library) and either Kapa2G Robust (Roche) or GoTaq (Promega) Polymerase. The PCR products were sequenced from the 5’ end by Sanger sequencing and resulting sequences were queried against the NCBI human RefSeq RNA database to identify the interacting human genes. The highest scoring match for each sequence is reported.

### Retest of Y2H interactions

Human gene inserts from AD plasmids were PCR-amplified and recloned into pOAD102 by *in vivo* homologous recombination in BK100 for genes that interacted with EBOV GP or BK100Nano (MATa ura3-52 ade2-101 trp1-901 leu2-3,112 his3-200 gal4Δ gal80Δ GAL2-Nluc LYS2::GAL1-HIS3 met2::GAL7-lacZ) for all other genes (35). Successful cloning was confirmed by PCR and sequencing. Y2H retest assays were performed by mating AD strains with BD strains expressing the viral protein that identified the interaction in Y2H library screens and selecting diploid yeast on SD-TL. For GP interaction, diploid yeast were plated on SD-TLUHA + 3-AT medium and incubated at 30° C. For all other interactions, diploid yeast were grown overnight in liquid SD-TL+A medium in 96-well plates at 30° C without agitation. The following morning, cultures were diluted 1:10 with fresh growth medium and incubated an additional three hours. Yeast were resuspended by pipetting and the OD_600_ was measured immediately using a Synergy 4 Multi-Detection Microplate Reader (BioTek Instruments, Inc., Winooski, VT, USA). Nano-Glo buffer and Nano-Glo substrate (furimazine, Promega, WI, USA) were reconstituted as per the manufacturer’s directions and added to samples at a 1:10 ratio (5 μL of reconstituted Nano-Glo substrate-buffer plus 50 μL of resuspended cell culture). Luminescence was measured 30 min after substrate addition using a Synergy 4 Multi-Detection Microplate Reader. Nanoluciferase values were divided by the OD_600_ value from the same well to correct for differences in growth. Each pair was tested in triplicate. Interactions were considered positive if the mean luciferase activity was greater than the mean luciferase activity plus five standard deviations from the negative control pairs (EBOV bait protein plus AD-GFP).

### Confidence scoring of Y2H interactions

Confidence scores were assigned to interactions based on rate at which they were positive in retest assays. Group 1 (high confidence, 369 interactions, 92% retest positive rate) included interactions that were positive in retest assays, that were identified more than once in one or more Y2H screens with the same bait, or that were identified one time with a gene whose relative abundance was less than 20 times the median gene abundance in the AD cDNA library. Group 2 (medium confidence, 32 interactions) included interactions that were identified only in screens with the with interferon or universal libraries, that were not retested, and that involved genes that were either not present in the liver or macrophage libraries or were present at a relative abundance of <50 times the median abundance. Group 3 (low confidence, 120 interactions) included interactions found once with a gene in the liver or macrophage libraries with a relative abundance greater than 10 (49 interactions, 44% retest positive); interactions that were negative in retest assays (29 interactions); interactions found only in screens of the interferon or universal libraries but that were present in either liver or macrophage libraries at a relative abundance of >50 (17 interactions); interactions involving human genes that interacted with >3 EBOV proteins (25 interactions).

### Analysis of enriched annotation terms

Gene enrichment analysis was performed using the Database for Annotation, Visualization, and Integrated Discovery (DAVID) v6.8 (37, 38). Lists of human genes identified in Y2H screens were analyzed against a background all genes detected by sequencing of the liver and macrophage cDNA libraries. Gene ontology terms with a false discovery rate (FDR) less than 0.05 were significant. Interaction networks were generated using Cytoscape v3.9.1.

### Plasmids and cloning

The full length PABPC1 ORF was cloned in frame with GFP in the NheI and BamHI sites at the 5’ end of GFP in plasmid pEGFP-N1 to generate pEGFP-N1-PABPC1 (Epoch Life Science, Inc.). PCDNA3-GFP1-9 T2A mCherry was a gift from Xiaokun Shu (Addgene plasmid # 124430; http://n2t.net/addgene:124430; RRID:Addgene_124430) (39). Plasmids pCDNA5-FRT-TO-EBOV-VP30-CT-GFP-B10, pCDNA5-FRT-TO-NT-GFP10-EBOV-VP40, pPABPC1-CT-GFP-B11, and pCDNA5-FRT-TO-NT-GFP-B11-TSG101 were described previously (40). Plasmid pGFP-B11-N1-RBBP6 was generated by digesting pPABPC1-N1-GFP-B11 with *Nhe*I and *Bam*HI and gel purified. The full length RBBP6 ORF which was PCR-amplified in two parts from pCAGGS-RBBP6 (a gift from Dr. Christopher Basler, (21)) using PCR primers 5’-TTTAGTGAACCGTCAGATCCGCTAGCCACCATGTCCTGTGTGCATTATAAATTTTC-3’ plus 5’-CTCTTTTGACTTTGTATTATCCTTCTG-3’ to amplify the 5’ half and primers 5’-CAGAAGGATAATACAAAGTCAAAAGA-3’ plus 5-GCTTTTCGCTACCTCCGCCTGGATCCACA GTGACAGATTTCACTTTTTGG-3’ to amplify the 3’ half. The PCR products were cleaned and inserted into digested pPABPC1-N1-GFP-B11 using the NEBuilder HiFi DNA Assembly (New England Biolabs #E2621) three fragment assembly protocol as described. Plasmid pGFP-B11-N1-RBBP6 Δ1 was generated by digesting pGFP-B11-N1-RBBP6 with *Bbv*CI and *Blp*I, gel purifying the larger vector band, and religating to generate an in-frame deletion of nucleotides 2653-5160 (amino acids 855-1720). Plasmid pGFP-B11-N1-RBBP6 Δ2 was generated by digesting pGFP-B11-N1-RBBP6 Δ1 with *Sal*I and *Bsg*I to delete nucleotides 1504-1878 (amino acids 502-626 containing the VP30 binding site) and gel purifying the larger vector band. The plasmid was religated using NEBuilder HiFi DNA Assembly master mix and oligo 5’-TGCTCGTCCAGGTGGTGGTACCATCCCAACAACACA-3’ to bridge the cut sites. All plasmids were verified by PCR and sequencing.

### Co-immunoprecipitations

HEK293T cells were seeded at a density of 7 x 10^5^ per well of a six-well culture plate. Cells were transfected the following day with 1 μg of each plasmid using Lipofectamine 2000 (ThermoFisher). After 24 h, cells were collected in a 1.5 mL sterile centrifuge tubes, washed three times with ice cold PBS, and lysed by adding 200 μL of lysis buffer (10 mM Tris-HCl pH 7.5, 150 mM NaCl, 0.5 mM EDTA, 0.5% Nonidet P40 (NP40)) supplemented with 1% protease inhibitor cocktail (Sigma Aldrich P8340) and 1 mM PMS-F. Cells were incubated 30 min on ice with occasional mixing. Cell debris was removed by centrifugation at 17,000 x g at 4° C for 10 min. Supernatants were removed and diluted with 300 μL of ice-cold lysis buffer without NP40. To assess the amount of protein in the input samples, 25 μL was removed from each sample, mixed with 2X SDS sample loading buffer and stored at -80° C. The remaining sample was mixed with 20 μL of GFP-Trap agarose beads (Chromotek, gta-100) equilibrated in lysis buffer lacking NP40. GFP capture was performed according to manufacturer protocol with the following modifications, except that capture time was reduced to 45 minutes to minimize the potential for proteolysis and wash volumes were increased to 1 mL. Samples were subjected to SDS-PAGE on SurePAGE Bis-Tris 4-12% gradient gels (GenScript) and transferred to nitrocellulose for western blotting. Blots were sequentially probed with rabbit polyclonal anti-Ebola VP30 antibodies (GeneTex, GTX134035) followed by IRDye 800CW goat anti-rabbit IgG secondary antibody (LI-COR, 926-32211); rabbit anti-GFP (Abcam, ab6556) followed by IRDye 680RD goat anti-rabbit IgG secondary antibody (LI-COR, 926-68071; and mouse anti-β-actin antibody (Sigma-Aldrich, A1978) followed by IRDye 680RD goat anti-mouse IgG secondary antibody (LI-COR, 925-68070).

### Split-GFP assays

Tripartite split-GFP assays were performed exactly as described by Mori et al (40). Briefly, HeLa cells were transfected in quadruplicate with plasmids expressing GFP1-9-T2A-mCherry, a GFP B11 fusion protein, and a GFP B11 fusion protein at a ratio of 5:5:1 with Lipofectamine 2000 (Invitrogen). At 48 h, cells were fixed with 4 % paraformaldehyde and stained with Hoechst 33342. and stored in the dark at 4° C until imaging. Fixed cells were stored at 4° c in the dark until imaging on a PerkinElmer Opera Phenix High Content Screening System. Nine fields per well were captured with a 10x/air NA 0.3 confocal lens and excitation/emission wavelengths (Exc/Em) of 375/435-480 (Hoechst33342), 488/500-550 (GFP), and 561/570-630 (mCherry). The mCherry and Hoechst were acquired simultaneously, whereas the EGFP channel was scanned independently.

After applying basic flatfield and brightfield corrections, image analysis was performed using Harmony High-Content Imaging and Analysis Software (version 4.9). First, nuclei were identified by mean intensity with area < 300 μm^2^, intensity < 6000 and roundness < 1.5 using method C. Second, transfected cells expressing mCherry were delineated using mean intensity with area > 100 μm^2^ using method B. Finally, total GFP spots in the transfected cell region of interest were enumerated based on intensity using method D. The nine fields from each well were merged without overlaps and the calculated output values were exported for statistical analysis in GraphPad Prism (GraphPad Software, San Diego, CA).

### Cultivation of Zaire EBOV

All EBOV experiments were performed under biosafety-level 4 (BSL4) conditions at the National Emerging Infectious Diseases Laboratories BSL4 suite in Boston, Massachusetts. Zaire EBOV was cultured on Vero E6 cells. Zaire EBOV was provided by Heinz Feldmann (Rocky Mountain Laboratories, Hamilton MT) and cultured on Vero E6 cells. Supernatants were collected 5 days post infection after development of cell-pathology effect and clarified via centrifugation. Virus was then concentrated over a 20% sucrose cushion via ultracentrifugation. Titer was calculated by serially diluting the virus on HeLa cells and staining for virus via immunofluorescence to the viral glycoprotein (antibody 4F3, IBT) in a focus-forming assay, 16 hours post infection.

### Generation of EBOV expressing VP30-FLAG

Recombinant EBOV (Genbank AF086833.2) expressing VP30 tagged with the FLAG epitope on the C-terminus was generated using circular polymerase extension reaction as previously described (41). Briefly, six overlapping EBOV fragments spanning the entire EBOV genome were amplified, purified, and assembled to generate a cDNA clone. Recombinant virus was recovered by transfecting HEK293T cells grown in a 12-well plate with 25 μL of CPER product, and support plasmids. Supernatants containing recombinant virus were collected four days post-transfection.

### Imaging viral protein and RNA in infected cells

HeLa cells were infected with VP30-FLAG recombinant EBOV at an MOI of 0.3 and fixed 18hpi in 10% neutral buffered formalin for at least 6 hours to inactivate EBOV. Samples were washed in PBS, permeabilized with 0.1% Triton X-100, and blocked in 3.5% BSA for 1 hour at room temperature. Primary antibodies were diluted in 3.5% BSA and incubated at 4° C overnight, washed with PBS, and then incubated with secondary antibodies diluted in 3.5 % BSA for 1 hour at room temperature. EBOV vRNA staining was performed as previously described except that the probes targeted NP vRNA (25). Following vRNA staining, cell nuclei were stained with Hoechst 33342 diluted in PBS. Z-stack images were acquired using a Nikon Spatial Array Confocal (NSPARC) Ti2 microscope using a 60 X oil immersion objective equipped with 640, 561, 488, and 405 lasers. Images were deconvolved using NIS Elements Deconvolution software (Nikon, Tokyo, Japan) and processed using Imaris software (Oxford Instruments, Concord, MA).

### Depletion of PABPC1 in HeLa cells

HeLa cells were seeded, incubated overnight and transfected with 5 nM PABPC1 or AllStars scramble siRNA (Qiagen) using Lipofectamine RNAiMax reagent (Invitrogen). At 24 hours post-transfection, cells were washed and incubated in DMEM + 10% FBS for an additional 24 hours. Samples were then lysed in RIPA Buffer (Boston Biosciences) with phosphoprotease inhibitor cocktail (Thermofisher) and boiled in 1% SDS for 10 minutes at 100⁰C. Samples were then analyzed via western blot to assess protein knockdown.

### Infection of PABPC1 depleted cells and assessment of EBOV replication

HeLa cells were transfected as described above. At 48 hours post transfection, cells were infected with EBOV at an MOI of 0.1. At 16 hours post infection, cells were either lysed with Trizol reagent or fixed in 10% neutral buffered formalin. RNA was collected from Trizol lysates via chloroform and isopropanol precipitation. Quantitative PCR of EBOV RNA was performed with primers and probes directed to positive-sense nucleoprotein RNA (sequences) using the Luna Universal One-Step RT-qPCR Kit. NP cycle values were normalized to beta-actin mRNA levels and transfected samples were normalized to the reagent-only treated control. RNA fluorescent-*in situ* hybridization (RNAFISH) to detect virus replication was adapted from (42) and performed as previously described (25). Cell nuclei were stained with Hoechst 33342 (Invitrogen) and samples were imaged on a Nikon Ti2 fluorescent microscope using a 10X objective. RNA expression was quantitated in ImageJ by measuring the area occupied by Cy5 signal and normalized to the area occupied by DAPI signal.

## Results

### Genome-wide Y2H screens for cellular proteins that bind to EBOV proteins

To identify human proteins that bind to EBOV proteins, we performed genome-wide yeast two-hybrid (Y2H) screens (Fig. 1). Multiple complementary strategies were employed to increase coverage and reduce false positives. First, we generated multiple bait constructs for each EBOV gene and expressed them as C- and N-terminal fusions to the Gal4 DNA binding domain (BD, also referred to as bait constructs). Using multiple constructs increases the likelihood of having at least one from each viral protein that is expressed well in yeast and that is functional in the Y2H assay. Large proteins such as L are poorly expressed as full length proteins, while smaller domains may be better expressed. Others may be toxic or inhibit growth when expressed as full length proteins, making it difficult to identify interaction partners. Screening with independent domains also enables us to narrow down the binding region. Second, bait constructs were cloned at 5’ and 3’ end of the Gal4 BD since N- and C-terminal fusions behave independently in the Y2H assay and the combination can be used to increase the number of interactions identified (43, 44). By including multiple overlapping constructs expressed as different fusions to the BD, redundancy is built into the process, which reduces false negatives (missed interactions) and increases confidence when the same interaction is found by partially overlapping constructs. Finally, we expressed the BD fusions using a strong yeast promoter to improve expression of difficult constructs and to increase the number of interactions (45).

**Fig. 1.**
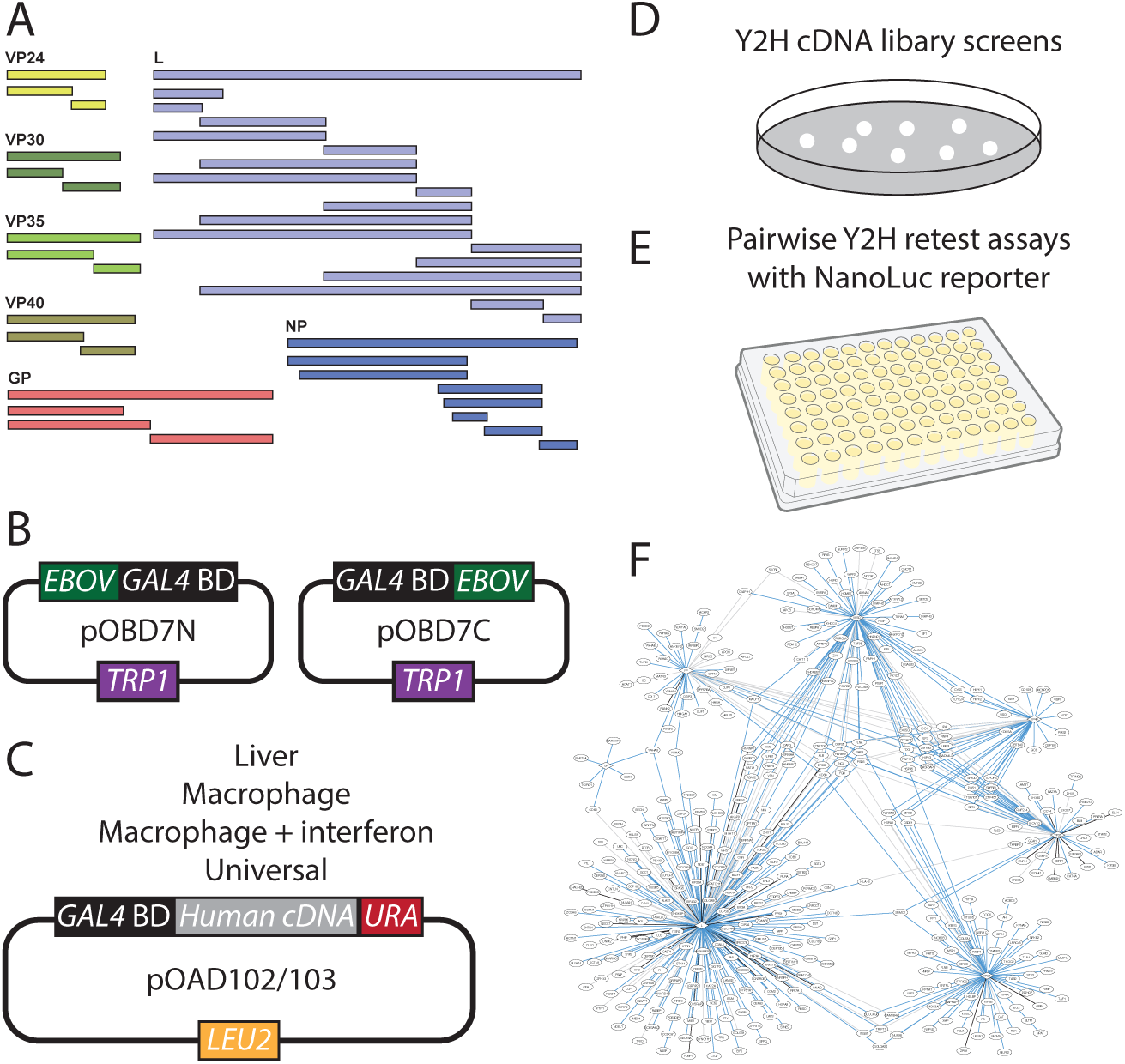
Overview of Y2H screens to identify EBOV-human protein-protein interactions. A-B. Full-length EBOV genes and gene fragments designed based on structural domains or homology to better characterized proteins were cloned at the 5’ and 3’ ends of the GAL4 DNA binding domain (BD) in pOBD7. C. Human cDNA libraries were generated from liver tissue, macrophages, interferon-treated macrophages, and a mixture of RNA from five human tissues (universal) in plasmids pOAD102 or pOAD103, which utilize *Saccharomyces cerevisiae URA3* or *Schizosaccharomyces pombe URA4* (*URA*) to select for cDNA fragments that are cloned in frame with the GAL4 activation domain (AD). D. EBOV BD constructs were screened against the cDNA libraries and positive colonies were selected under stringent growth conditions. E. A subset of genes from the cDNA library screens were recloned into pOAD102 and retested for interaction in BK100Nano, which uses NanoLuciferase as the quantitative Y2H reporter gene. F. A network of EBOV-human protein interactions with confidence scores was generated from cDNA library screens and retest assays.

Each viral gene was expressed as a full-length clone and as individual protein domains that were designed based on published structural data (Table S1, Fig. S1). In cases where structural information was not available, fragments were constructed based on alignments to related viral proteins with known structures and/or to other family members. In the latter case, regions of low sequence conservations were used as break points between fragments. In the case of L, domain fragments were based primarily on the alignment of EBOV L to the vesicular stomatitis virus (VSV) L protein, whose structure has been resolved by cryo-electron microscopy (46). GP constructs were based on the domain structure of the complete glycoprotein and truncated soluble GP (47). In total, 100 constructs were generated for the 7 EBOV genes. Each bait was tested for self-activation and the concentration of 3-AT needed to inhibit growth on medium lacking histidine was used for library screens.

Activation domain libraries were constructed in the Y2H plasmids pOAD102 and pOAD103. These plasmids encode auxotrophic markers (*Schizosacharomyces pombe* URA4 and *S. cerevisiae* URA3, respectively) downstream of the Gal4 activation domain (AD) (33, 34). In the parental plasmid, the auxotrophic marker is out of frame with the activation domain, so yeast containing this plasmid are unable to grow on media lacking uracil. However, upon introduction of a gene fragment that is in frame with both the AD and Sp-URA4/Sc-URA3, the auxotrophic marker is produced as a fusion protein and the yeast gain the ability to grow on media lacking uracil. Thus, this system enables gene fragments that are not in frame with the AD or that are not expressed to be excluded from the AD library. If the fragments were randomly generated exclusively from ORFs, one out of nine (11%) would be expected to grow on medium lacking leucine and uracil. In practice, ∼1-2 % of the clones that grow on medium lacking leucine are also able to grow on medium lacking uracil, the difference likely being due to the presence of stop codons in 5’ and 3’ UTRs and poor expression of some gene fragments. The practical effect of this approach is that the AD libraries are smaller than traditional libraries, making them easier to screen comprehensively, and have excluded clones in non-expressed reading frames, which generate short peptides that are a major source of false positives from cDNA library screens.

EBOV baits were screened against an existing AD library prepared from human liver RNA (36) and three new libraries generated from RNA isolated from human macrophages, human macrophages exposed to recombinant interferon-β to enrich for interferon-stimulated genes (ISGs) and a mixture of total RNA from five human tissues meant to represent expression of >90% of predicted human genes (referred to as liver, macrophage, interferon and universal libraries, respectively). In our initial screens, we found that the liver and macrophage libraries yielded the most interactions, so we focused our efforts on those libraries for the remainder of the study.

To assess the coverage of the liver and macrophage libraries, we amplified the human gene inserts from the libraries and sequenced them. The reads were mapped to a total 12,402 and 11,517 unique Ensembl IDs for the liver and macrophage cDNA libraries, respectively, of which 12,242 and 11,396 were protein coding (Table S2). Surprisingly, the libraries had extensive overlap, with 10,680 Ensembl IDs detected in both libraries, including 10,596 annotated as protein coding. 13,237 unique Ensembl IDs were found in the union of both libraries, representing 13,042 protein coding genes or about 64% of the protein coding genes in Ensembl (48). The overlap in the libraries could be due in part to the fact that the liver library was generated from RNA isolated from human tissue. The liver is a mixture of cell types, primarily hepatocytes, but also stellate, endothelial, and Kupffer cells, the latter of which are a type of resident macrophage (49, 50). The inclusion of non-protein coding sequences was unexpected, but could be due to open reading frames in non-coding sequences that enable read-through from the AD to the auxotrophic marker, sequences that act as cryptic promoters that allow the auxotrophic marker to be expressed in the absence of a fusion to the AD and in-frame human gene fragment, or sequences that are recognized as splice sites that allow the auxotrophic marker to be expressed as a fusion to the AD. Although there is extensive overlap in the genes in the cDNA libraries, the genes are represented by different fragments (libraries were constructed by random priming) and are present at different levels. The liver library was dominated by albumin, which accounted for nearly a third of the reads. The ten most abundant clones (albumin, plus fibrinogen alpha and beta, apolipoproteins B and H, transferrin, serpin A1) represented 49% of the clones in the library. In contrast, the ten most common genes in the macrophage library accounted for 9% of the clones, with ubiquitin C (UBC) being the most abundant.

From more than 200 Y2H library screens, we identified 3437 pairs of EBOV and human proteins representing 521 unique interactions (Table S3). Despite significant overlap between the libraries, the macrophage library yielded substantially more interactions than either the interferon or liver libraries (Table 1). Twice as many interactions were identified exclusively in the macrophage library than in the interferon library and 33% more were identified in the macrophage library than in the liver library.

**Table 1.** Summary of EBOV yeast two-hybrid screens.

| Library | Total pairs<br>identified | Unique interactions | Unique to library |
| --- | --- | --- | --- |
| Interferon | 994 | 168 | 86 |
| Liver | 838 | 209 | 135 |
| Macrophage | 1485 | 286 | 182 |
| Universal | 120 | 6 | 2 |
| Total | 3437 | 521 | - |

To assess the quality of our Y2H screens, we retested a subset of the interactions. The human AD inserts from 229 genes were PCR amplified, cloned into the parental AD plasmid by homologous recombination in the yeast strain BK100 (for GP-interacting genes) or BK100Nano (all others) and confirmed by sequencing (35). In general, AD constructs were tested for interactions with any fragment from the cognate viral gene that yielded interactions from the Y2H library screens (e.g., if VP40 fragment 1 identified an interaction with human gene X, it was retested for interactions with VP40 fragment 1 plus any other VP40 construct that yielded an interaction in the Y2H library screens). Strains expressing bait and prey constructs were mated, propagated on medium lacking tryptophan and leucine to select diploid yeast containing both plasmids, grown overnight in liquid YPDA, and assayed for luciferase activity. As a control, yeast expressing the bait construct were mated with a control strain in which GFP was inserted into the AD plasmid. At least eight replicates of the GFP negative control were included on each plate. Interactions were considered positive if the luciferase activity was greater than the mean GFP activity plus five standard deviations (Fig. S2, Table S4).

We considered an interaction to be positive in the retest assays if any construct from the viral gene that identified the interaction in Y2H screens was positive. Overall, 87% of the interactions tested were confirmed in at least one Y2H screen, whereas 77% of the specific viral protein construct-human protein interactions were positive. To determine which factors contributed to positive and negative retest results, we compared the retest results to parameters from the Y2H screens (Fig. 2A). Interactions that were found in multiple independent Y2H screens, either with the same construct in different libraries or using different constructs with the same library, retested at a higher rate than those that were not. Conversely, interactions that were found a single time in one screen were the least likely to be positive in retests. Nonetheless, 72% of the interactions in this group retested positive.

**Fig. 2.**
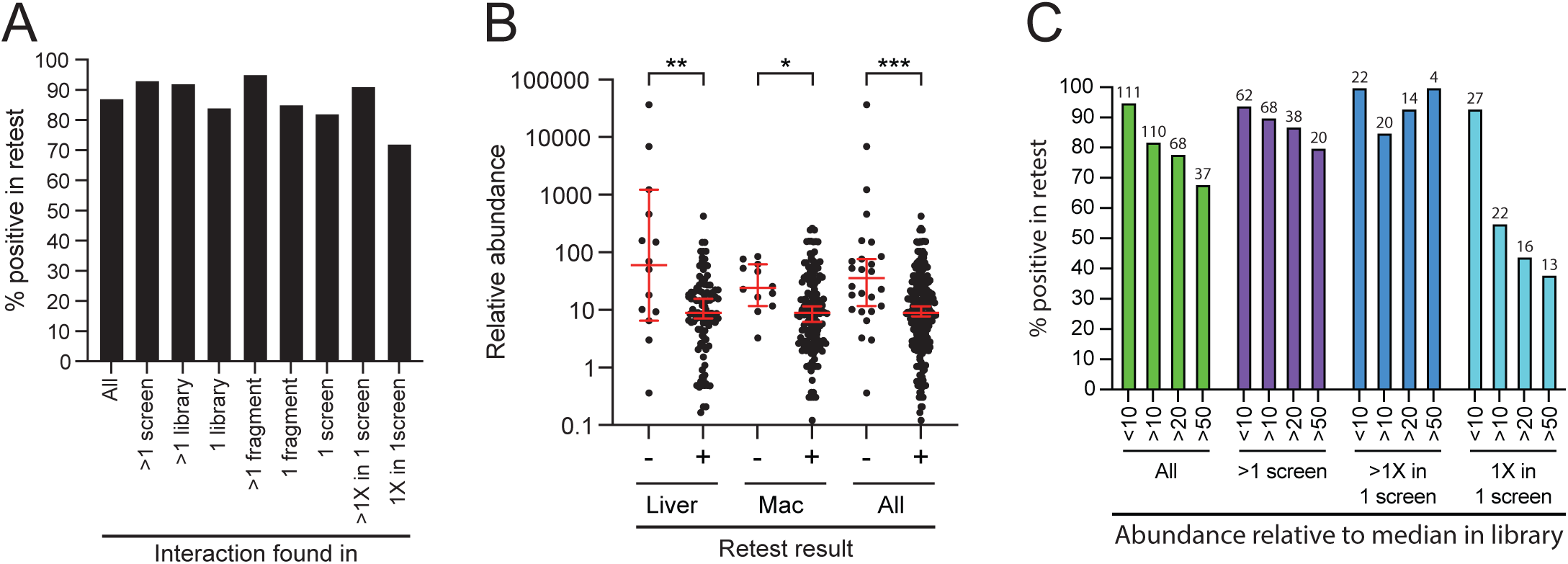
Correlation of EBOV-human gene retest success with Y2H screening parameters. A. Positive retest rates were graphed as a function of how often the interaction was identified in Y2H cDNA library screens. Interactions found multiple times in the same or in independent screens retest positive at higher rates. B. Graph shows abundance of human genes from interactions that were negative (-) or positive (+) in Y2H retest assays. Abundance is normalized to the median abundance of all genes in the library. Mean normalized abundance and standard deviation is indicated by red lines. Interactions that were negative in retest assays tended to be more abundantly represented in the libraries. C. Graph shows percentage of retest reactions that were positive as a function of abundance in the cDNA libraries relative to the median abundance of all genes. The four sets of bars show all interactions (green bars), interactions found in more than one independent screen (purple bars), interactions found more than once in one screen (blue bars) and interactions found one time in one screen (cyan bars). Numbers at the top of each bars indicate the number of interactions represented. Interactions found once in one screen with human genes that were more than ten times as abundant in the cDNA libraries as the median gene were the least likely to be positive in retest assays.

We also assessed whether the abundance of the human genes in the Y2H libraries was correlated with the likelihood of a positive result in the retest assay (Fig. 2B-C). To simplify presentation of the data, the abundance of each gene in the library was normalized to the median abundance of all genes in the library. The median abundance of genes identified as EBOV partners in Y2H screens of the liver and macrophage libraries was 9-10 times the median abundance of all genes in library. Genes in interactions that were positive in Y2H retest assays had a similar median abundance to genes identified in cDNA library screens. However, genes from interactions that were negative in retest assays were 7- and 2.7-times more abundant in the liver and macrophage libraries, respectively, than genes from interactions that tested positive. This suggests that more abundant genes were more likely to lead to false-positive interactions. Based on these observations, we assigned the remaining interactions that were not retested to three groups: 1 (high confidence, 369 interactions, 92% retest positive rate), 2 (medium confidence, 32 interactions), and 3 (low confidence, 120 interactions) (Table S5, Fig. S3).

### Y2H screening identifies novel and previously reported proteins implicated in EBOV infection

To assess the quality of the Y2H data set, we compared the set of interactions to published reports of EBOV protein-protein and functional interactions. Forty-six EBOV-human protein interactions were previously reported, and another 11 proteins from the Y2H screen were detected in EBOV virions (Fig. S4, Supplementary Tables 6 and 7). Comparing the genes identified in the Y2H screen to a recent genome-wide optical pooled screen of CRISPR knockouts revealed 61 genes that significantly affected EBOV replication or c-Jun nuclear localization (Fig. 3, Table S8) (51). In general, proteins that were required for EBOV replication had similar effects on EBOV RNA and protein production, with TUFM having the largest impact (Fig. 3A). TUFM, a mitochondrial elongation factor, interacted with NP in this study and was also reported as a NP-binding partner in co-immunoprecipitation and proximity-dependent biotinylation experiments (28, 52, 53). Knockouts of two genes identified in the Y2H screen (TLN1 and ITGB1) caused an increase in EBOV RNA and protein, whereas knockout of CSDE1 increased EBOV RNA without affecting VP35 protein levels (51); all three are high confidence partners of VP24. In contrast, two genes from the EBOV Y2H screen (PABPC1 and PDCD2) caused disproportionate reductions in EBOV VP35 RNA levels relative to VP35 protein when they were knocked out (51). Similar patterns were observed with c-Jun nuclear localization, with most knockouts reducing nuclear c-Jun localization proportionally to VP35 protein levels, suggesting that reduced c-Jun trafficking was due to lower EBOV replication (Fig. 3B). However, deletions of eight gene encoding proteins that interacted with EBOV proteins in this study significantly increased c-Jun nuclear localization without affecting EBOV replication. Together, these studies revealed extensive overlap between the hits from the EBOV Y2H screen and published reports and provide potential mechanistic explanations for the effects of gene knockouts in the genome-wide optical pooled screen (51).

**Fig. 3.**
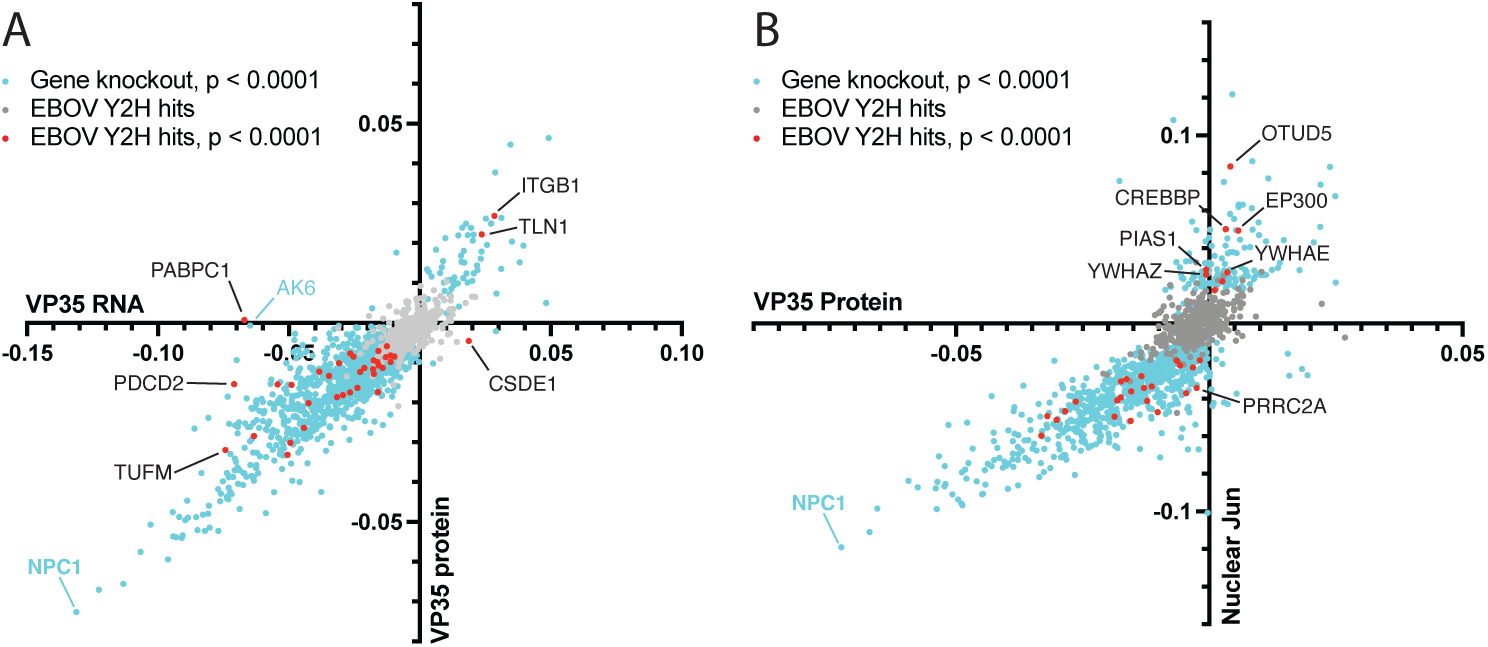
Integration of host factors from EBOV Y2H and genome-wide CRISPR-Cas9 knockout optical pooled screens (51). VP35 RNA and protein abundance from the Carlson et al. primary screen, Table S1, are replotted in A and B with Y2H screen hits highlighted (51). A. VP35 Protein Median Cell Mean cumulative delta AUC was plotted as a function of VP35 RNA FISH Median Cell Mean cumulative delta AUC. Graph shows human genes that had a significant effect on VP35 RNA or protein levels (p<0.0001, light blue dots), human genes identified in the EBOV Y2H screen that did not have a significant effect on EBOV RNA or proteins levels in the EBOV OPS (p>0.0001, gray dots), and human genes identified in the EBOV Y2H screen that had a significant effect on EBOV RNA or proteins levels in the EBOV OPS (p<0.0001, red dots). B. Jun Median Nuclear Mean cumulative delta AUC (y-axis) was plated as a function of VP35 Protein Median Cell Mean cumulative delta AUC (x-axis). Graph shows human genes that had a significant effect on nuclear Jun levels (p<0.0001, light blue dots), human genes identified in the EBOV Y2H screen that did not have a significant effect on nuclear Jun levels in the EBOV OPS (p>0.0001, gray dots), and human genes identified in the EBOV Y2H screen that had a significant effect on nuclear Jun levels in the EBOV OPS (p<0.0001, red dots).

### Enriched functional and structural features in the EBOV-human protein interaction network

To identify enriched terms in the EBOV Y2H data set, human genes from high and medium confidence interactions were submitted to the DAVID database using the set of genes identified in the liver and macrophage libraries as background. Among the entire data set, regulation of transcription, mRNA processing and sumoylation were the most enriched terms (Fig. 4 and Table S9). Similar to flavivirus RNA-dependent RNA polymerases, L binding partners were enriched for cellular proteins that localized to the centrosome (36, 54). RNA binding proteins were prominent among the partners of L and VP30, whereas dsRNA binding proteins were enriched among the partners of VP35. VP35 is known to bind dsRNA (55–61); it is not known if the interactions detected between VP35 and cellular dsRNA binding proteins is direct or mediated by dsRNA. Proteins involved in sumoylation were prominent partners of VP35 and VP40 (62, 63). Transcription related proteins were enriched among the partners of L and VP40. L, NP, VP24, VP35, and VP40 targeted zinc finger proteins, though they bound to different cellular proteins and with different types of zinc fingers. VP30 partners were enriched in RNA recognition motif (RRM) domains. NP partners were enriched for 14-3-3 (YWHAB, YWHAE, YWHAG, YWHAH, YWHAQ, and YWHAZ) and the MYND (MYeloid translocation protein 8, Nervy and DEAF-1, (64)) domain proteins (ANKMY2, PDCD2, and SMYD2). All 14-3-3 domain proteins and two of three MYND domain proteins bound to an NP fragment containing the large intrinsically disordered region. Hower, no canonical 14-3-3 phosphopeptide binding sites are present in this region of NP, suggesting that 14-3-3 proteins bind to a noncanonical sequence (65, 66).

**Fig. 4.**
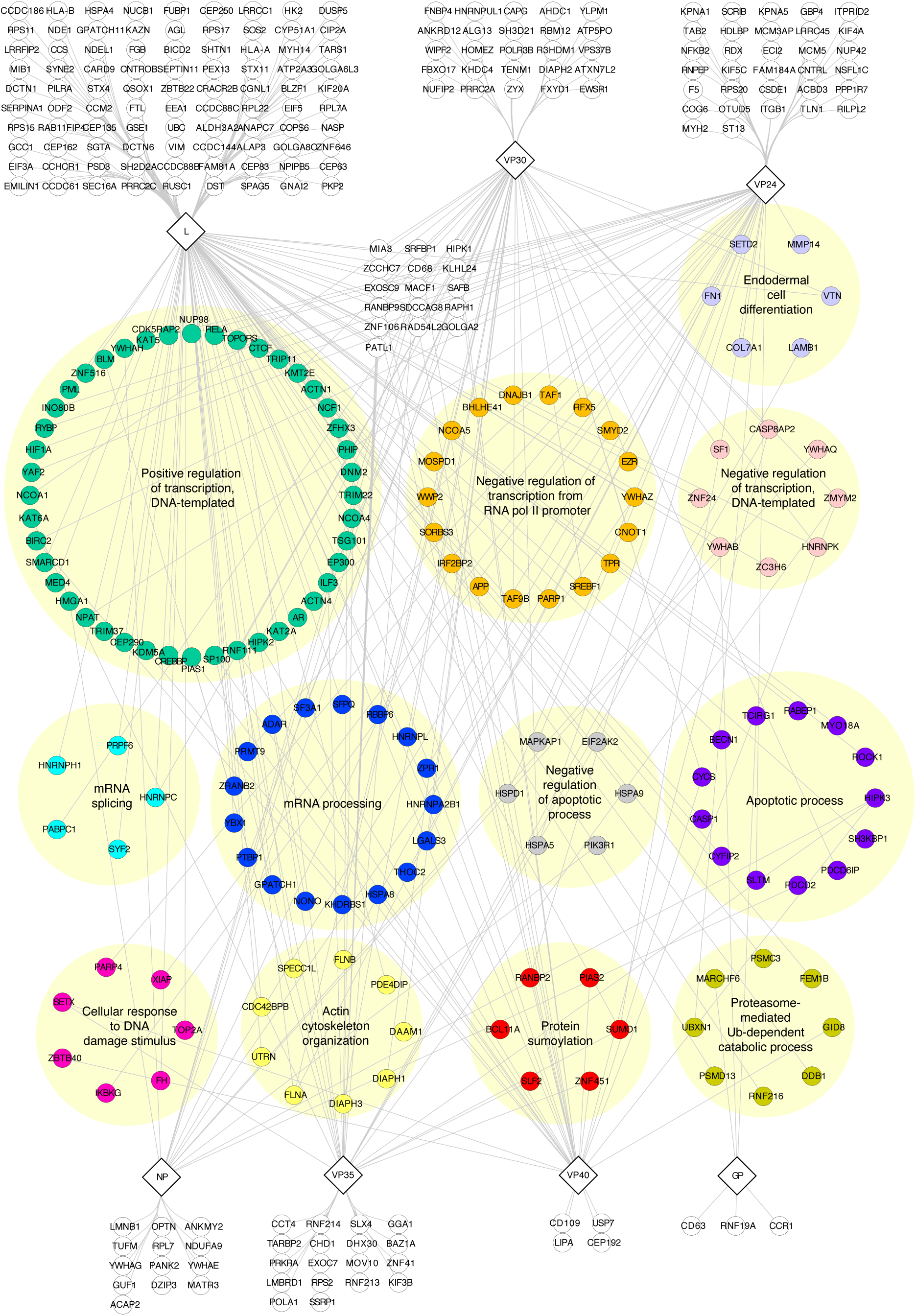
Enriched features in EBOV-human network. Network of high and medium confidence EBOV-human protein-protein interactions identified in the EBOV Y2H screens. Human proteins (circles) are grouped according to their enriched GO Biological Process terms. EBOV proteins are shown as diamonds. The full list of enriched terms for the entire Y2H data set and individual proteins is in Table S9.

### EBOV NP interacts with MYND domain proteins

The MYND domain is a C6HC zinc finger found in diverse proteins that mediates protein-protein interactions via binding to PxLxP, PxLE and PPxLI motifs (67–69). Three MYND domain proteins interacted with NP in this study and two others, EGLN1 and SMYD3, were previously reported to bind to NP (Fig. 5A) (24, 28, 52). The MYND domain proteins that interacted with NP have no homology to each other outside of the MYND domain, suggesting that NP binding is via the MYND domain. A putative MYND binding site is found at amino acids 583-586 in Zaire EBOV NP (SSL**<u>P</u>**P**<u>LE</u>**) between the serine/threonine protein phosphatase 2A (PP2A) binding site and the VP30 binding site within the NP intrinsically disordered region (IDR; residues 424-651) (70). This sequence closely resembles the known binding sites of ANKMY2 and EGLN1 in FKBP8 (SEL**<u>P</u>**P**<u>LE</u>**) and PTGES3 (also known as P23; KM**<u>P</u>**D**<u>LE</u>**) (Fig. 5B) (69, 71–73). A prior analysis of the interaction of SMYD3 with NP is consistent with binding being mediated by the SMYD3 MYND domain (24). Furthermore, mutating the P and L of the PxLE site in NP disrupted NP binding to SMYD3 but did not affect recruitment of PP2A-B56 or VP30 (24).

**Fig. 5.**
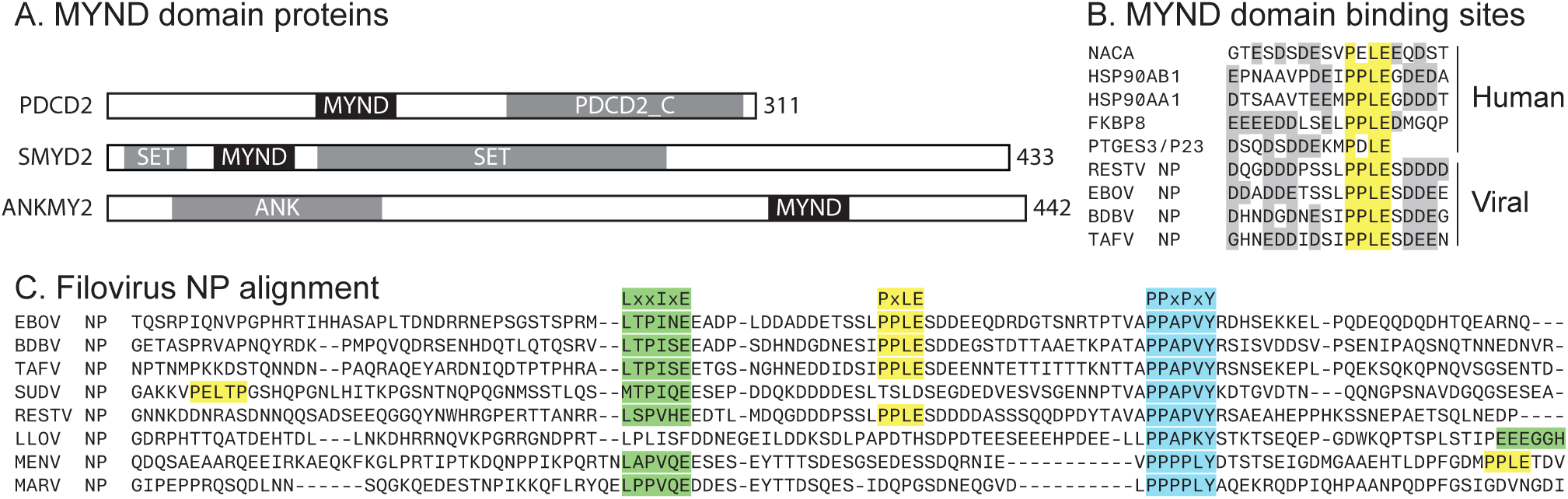
EBOV NP binds to MYND domain proteins. A. Schematic representation of human MYND domain proteins identified in the EBOV Y2H screens. MYND (MYeloid translocation protein 8, Nervy and DEAF-1) domains are represented by black bars and other domains are indicated with gray bars. Numbers indicate amino acid positions. PDCD2_C, Programmed cell death protein 2, C-terminal putative domain; SET, Suppressor of variegation 3-9 (Su(var)3-9), Enhancer of zeste (E(z)), and Trithorax (Trx) domain; ANK, ankyrin repeats. B. Alignment of human and predicted filovirus MYND domain sites. Conserved residues in the MYND domain binding site are highlighted in yellow. Negatively charged residues flanking the MYND domain binding site are shaded in gray. C. Alignments of filovirus NP proteins. Protein phosphatase 2A (PP2A), MYND domain, and VP30 binding sites are highlighted in green, yellow and blue, respectively.

The putative MYND domain binding site is mostly conserved among members of the genus *Orthoebolavirus* (Fig. 5C) but absent in Marburg virus (MARV). It diverges in Sudan virus (SUDV) by virtue of a T in place of the critical first P (24), which suggests that SUDV NP likely does not bind to MYND domains at this site. However, SUDV has an exact PxLxP motif at position 527-531 (**<u>P</u>**E**<u>L</u>**T**<u>P</u>**), suggesting SUDV has evolved an independent MYND binding site not found in other *Filoviridae*. Like MARV and SUDV, the bat filoviruses lack the PxLE sequence at this site, though Mengla virus has a PPLE motif in the NP IDR (589-592, **P**P**LE**TD) downstream of the VP30 binding site. A similar rearrangement of the order the binding sites was also noted for Lloviu virus, in which the PP2A-B56 binding site is downstream from the VP30 binding site (20).

### Domain mapping of interactions

Use of EBOV gene fragments enabled us to define which regions or domains of EBOV proteins mediated the interactions (Fig. 6). Most interactions with NP mapped to internal fragments from the intrinsically disordered region, which contains short linear sequences that interact with VP30 and the cellular protein phosphatase (20, 74, 75). Similarly, most interactions with VP30 map to the C-terminal domain, which binds to both NP and many cellular proteins through the same binding site. Interactions with L primarily mapped to constructs from the C-terminal region, which include the connector, methyltransferase and C-terminal domains. In contrast, few interactions mapped to L fragments corresponding to the RdRp. Since the RNA polymerase must undergo extensive conformational changes during transcription and genome replication, it may be that interactions with host factors are not compatible with enzymatic activity.

**Fig. 6.**
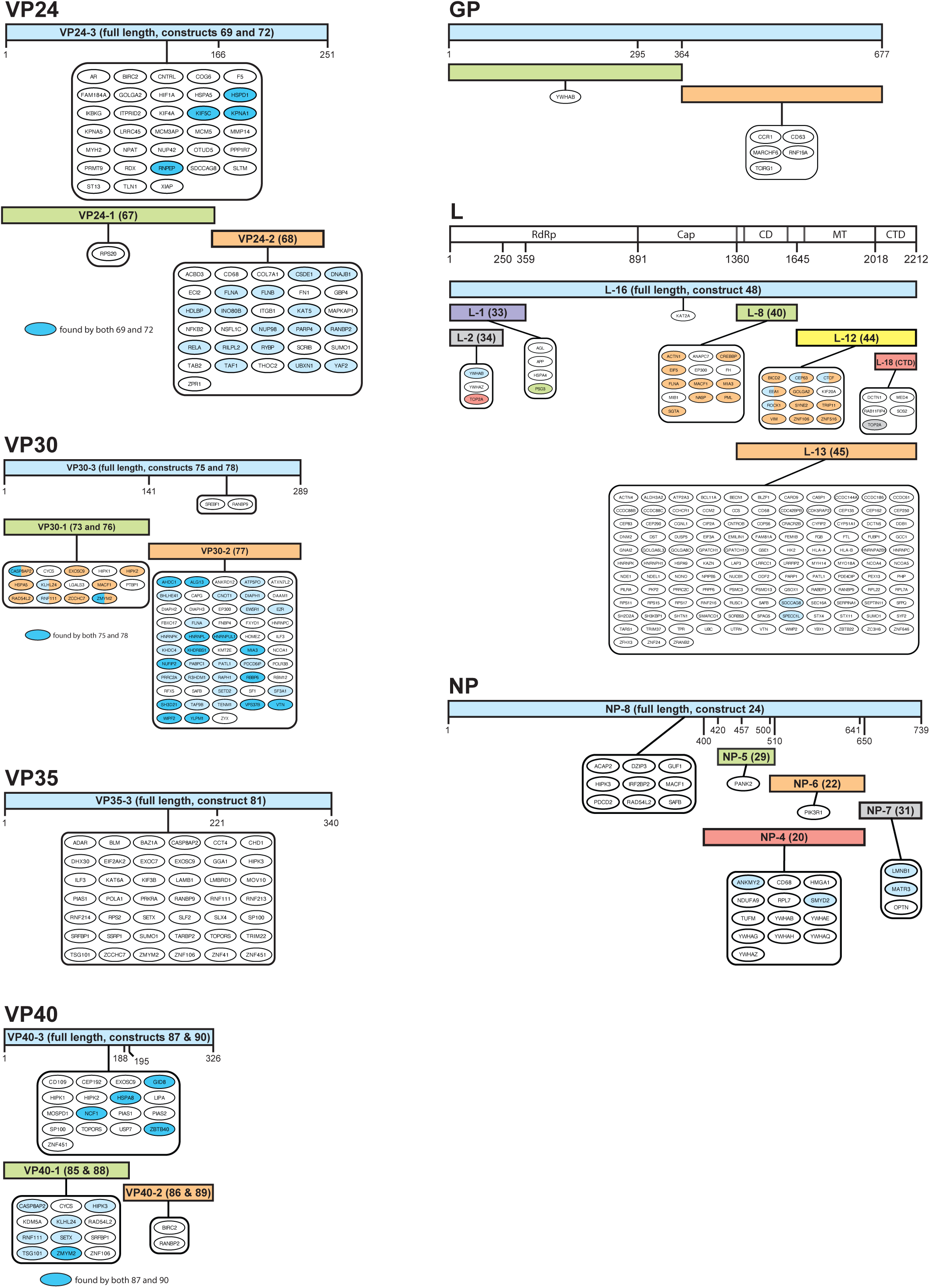
EBOV-human protein interactions mapped to viral protein domains. Human proteins that interacted with EBOV proteins are shown associated with the smallest protein domain that identified the interaction. Human proteins that were identified by other larger domains are shaded as indicated in the figure.

Because we used cDNA fragments instead of full length ORFs for our AD libraries, we were able to obtain information about which domains bound to the EBOV proteins in many interactions. In some cases, partially overlapping clones from the Y2H screens enabled more precise mapping of the binding site on the cellular protein (Fig. 7). In the best example, two partially overlapping clones from RBBP6 revealed a shared 20-amino acid segment (residues 552-571) that included the known VP30 binding site (560-PPPPLY-565) (21). Using similar overlapping clones, we hypothesize the VP30 binding site on NUFIP2 is located within amino acids 346-377, the binding site for L on SDCCAG8 is between amino acids 104 and 147, and the binding site for VP35 on SETX is in a 19 amino acid sequence from 1009-1027.

**Fig. 7.**
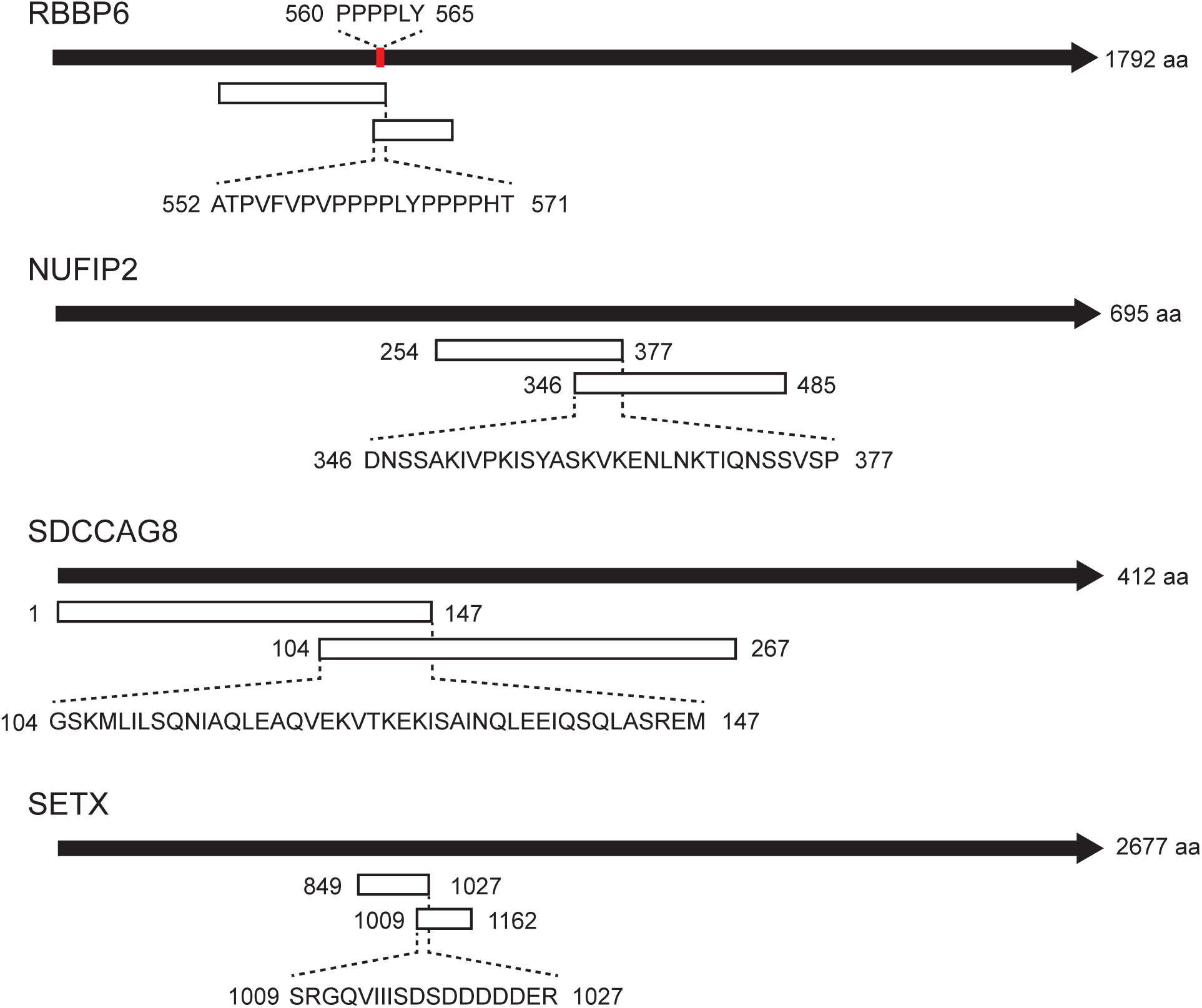
Identification of predicted EBOV protein binding regions using partially overlapping clones from Y2H screens. Black bars indicate human proteins. White bars represent fragments from AD clones identified in Y2H screens. Red bar in RBBP6 indicates the VP30 binding site. Numbers indicate amino acid positions. Amino acid sequences from partially overlapping AD fragments, representing predicted binding regions for viral proteins, are shown in brackets.

### VP30 interacts with proline-rich and non-proline-rich partners

VP30 binds to NP and host cell proteins containing the sequence motifs PPxPxY or PxPPPPxY via a site in the C-terminal domain (21, 22, 74, 75). To determine if the VP30-binding partners identified in the Y2H screens contained similar motifs, we analyzed the full-length sequences with the Multiple EM for Motif Elicitation program (MEME) (76). MEME correctly identified known VP30-binding sites in RBBP6, HNRNPL, and HNRNPUL1, as well as consensus VP30-binding motifs in EZR (PPPPPPPVY) and PRRC2A (PVPPPPPY) (Table S10). Three proteins have proline rich sequences followed by tyrosine that diverge slightly from the consensus motifs (ALG13 (PPPPLPPPPY), WIPF2 (LAPPPPPY and APPPPPPY), and YLPM1 (PPPQPPPSY)). Thirty other proteins contained stretches of multiple proline residues without a tyrosine (Fig. S5), suggesting that VP30 has the potential to bind to a wide array of proline-rich sequences, at least in the Y2H assay. The remaining VP30-binding proteins lacked motifs with consecutive prolines. In some case these proteins were frequently identified and had nanoluciferase activity similar to validated VP30 interactors such RBBP6 and HNRNPUL1. For example, NUFIP2 was the most commonly identified VP30 partner in Y2H library screens and yielded nanoluciferase reporter activity comparable to known interactors. VTN was identified as a VP30 partner a single time in library screens but had the highest nanoluciferase activity.

To determine if the VP30 interactors identified in the Y2H screens bound to the same site as NP and cellular proteins that contain PPxPxY or PxPPPPxY motifs, we tested the effect of mutations in the NP/RBBP6 binding site (74, 75). Fifty cellular genes that interacted with the C-terminal domain of VP30 were tested in pairwise Y2H assays with nanoluciferase as the reporter gene (Fig. 8, Fig. S6, Table S11). W230A had the most dramatic effect, eliminating binding to proteins that lacked poly-proline sequence or that contained poly-proline sequences or consensus VP30 binding motifs. D202A had a similarly consistent effect, reducing but not eliminating binding to nearly all partners. E197A and R213A, which disrupt binding to NP, had variable effects. E197A reduced binding to about half of the proteins that lacked poly-proline sequences but had no effect or even increased binding to proteins with poly-proline sequences or consensus VP30 binding motifs. In contrast, R213A had little to no effect on binding to proteins that lacked poly-proline sequences but substantially reduced binding to proteins with poly-proline sequences or consensus VP30 binding motifs. However, in two cases, HNRNPL and EWSR1, R213A significantly increased binding; pairwise BlastP analysis did not reveal significant homology between HNRNPL and EWSR1. Together these data suggest that proteins containing or lacking poly-proline sequences bind to the same site on VP30 but likely make different contacts with VP30 residues within the binding pocket.

**Fig. 8.**
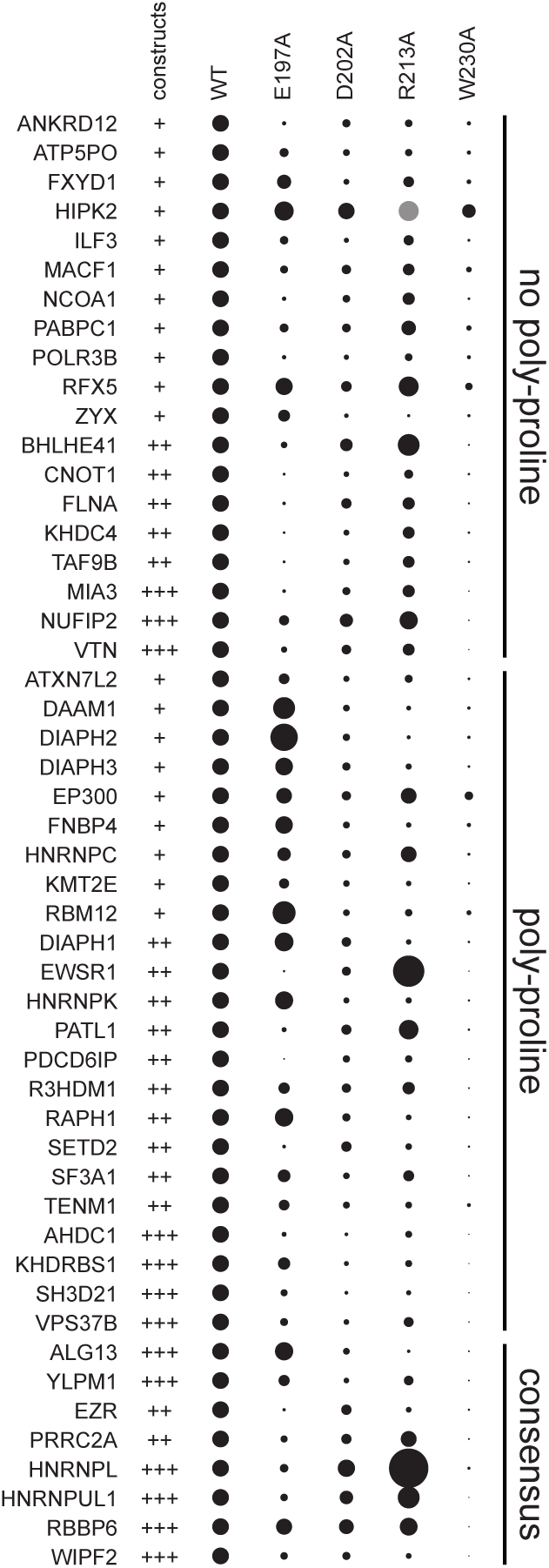
Effect of mutations in VP30 C-terminal domain on binding to human proteins. Wildtype VP30 C-terminal domain and VP30 mutants E197A, D202A, R213A, and W230A were tested for binding to fragments from the indicated human genes in pairwise Y2H assays using nanoluciferase reporter gene activity as an indicator of interaction strength. Nanoluciferase activities for each gene were normalized to wildtype. Dot size corresponds to relative nanoluciferase activity. Wildtype VP30 constructs that interacted with each human gene are indicated in the first column (+, VP30 C-terminal domain fused that the N-terminus of BD; ++, VP30 C-terminal domain and VP30 full length fused that the N-terminus of BD; +++, VP30 C-terminal domain, VP30 full length fused that the N-terminus of BD, and VP30 full length fused that the C-terminus of BD.

### PABPC1 binds to EBOV VP30

To assess the functional significance of VP30 binding proteins that lack a poly-proline motif, we focused on poly(A) binding protein cytoplasmic 1 (PABPC1, also frequently referred to as PABP). PABPC1 was independently identified as a partner of EBOV VP30 by proximity dependent biotinylation and is a common target for many viruses (52, 77). The 636 amino acid protein consists of four copies of the RNA recognition motif (RRM) at the N-terminus, a flexible proline-rich connector domain, and a conserved, structured domain at the C-terminus (PABC, Fig. 9A) (78). The proline rich linker, as its name implies, is rich in proline residues, but lacks stretches of consecutive prolines present in consensus VP30-binding motifs or in the poly-proline motifs identified in VP30 partners by MEME. Based on partially overlapping clones from the VP30 Y2H library screens, the likely binding region is located within residues 398-532, which includes parts of the proline-rich linker and the PABC domain. Since PABPC1 interacts with the VP30 C-terminal domain, which lacks the RNA binding sequences in the N-terminal domain, the interaction between is unlikely to be mediated by RNA.

**Fig. 9.**
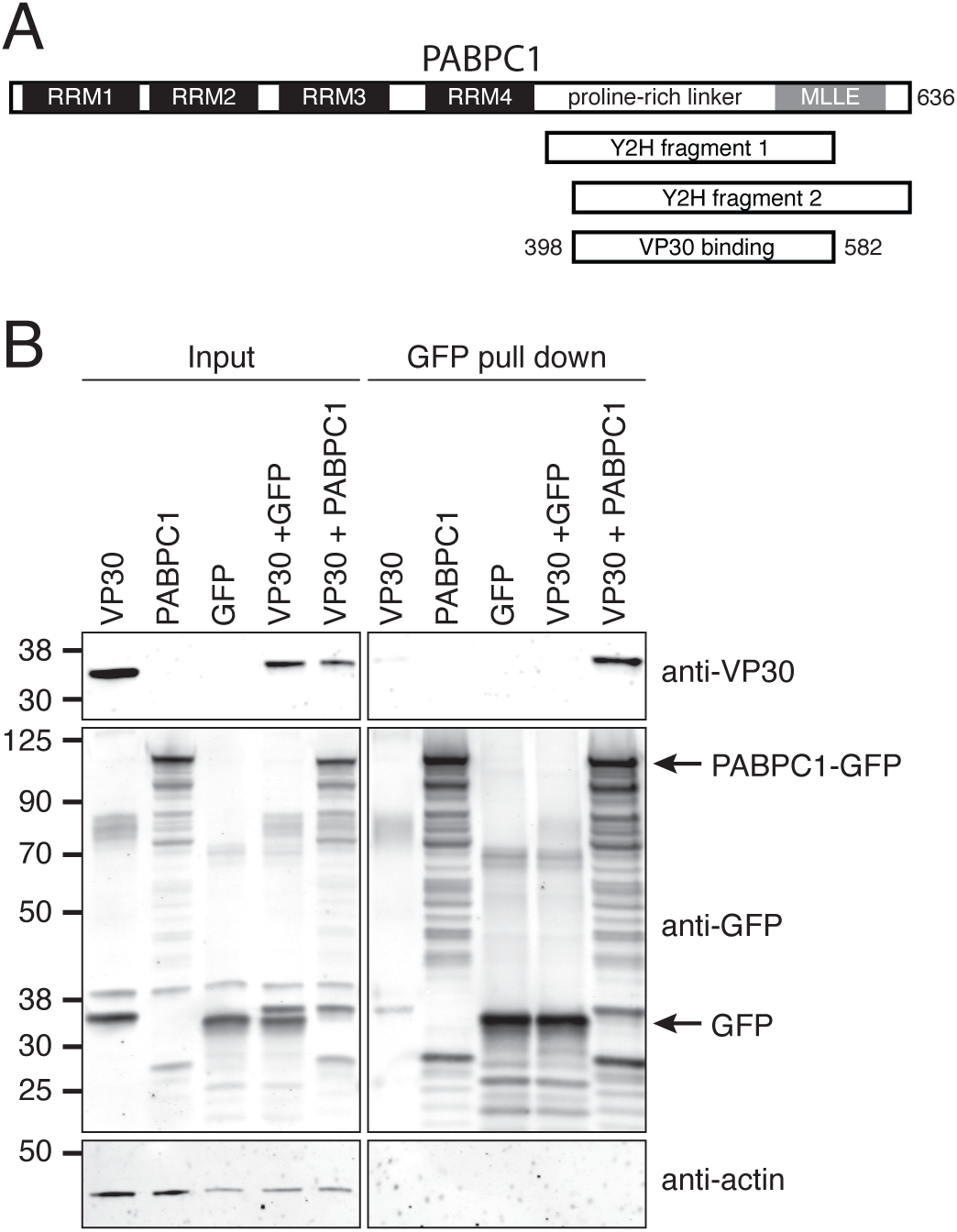
Co-purification of EBOV VP30 with PABPC1. A. Diagram of PABPC1 showing RRM (black bars) and PABP (gray bar) domains. Y2H fragments from VP30 screens and the predicted VP30 binding region are shown below. B. HEK293 cells were transfected with VP30, GFP and PABPC1-GFP expression plasmids and lysed. PABPC1 was precipitated with anti-GFP nanobodies and the precipitants were subjected to SDS-PAGE followed by western blotting with VP30, GFP, and actin antibodies. Input and precipitated samples are shown on the left and right, respectively. Actin loading control is shown in the bottom panel. Molecular weight markers in kDa are shown at left.

To determine if PABPC1 interacted with VP30 in mammalian cells, we performed co-immunoprecipitations with PABPC1 fused to GFP. HEK293 cells were co-transfected with plasmids expressing wildtype VP30 and either GFP or PABPC1-GFP. Immunoprecipitations with an anti-GFP nanobody was then performed on soluble lysates. Both GFP and PABPC1-GFP were efficiently precipitated with anti-GFP nanobodies, but VP30 co-purified only with PABPC1-GFP, indicating that PABPC1 associated with VP30 (Fig. 9B).

As an independent approach to confirm the interaction, we implemented a tripartite split-GFP assay in which putative interacting proteins are tagged with the 10^th^ and 11^th^ beta strands of GFP and co-expressed with GFP_1-9_. Complex formation by GFP_10_- and GFP_11_-tagged proteins enables complementation of GFP1-9, resulting in green fluorescence. To establish the tripartite split GFP system with VP30, we first evaluated the interaction of VP30 with RBBP6 (Fig. 10A). C-terminally GFP_11_-tagged full length RBBP6 was unable to complement GFP_1-9_ when co-expressed with VP30-GFP_10_. Due to its large size (1792 amino acids), we hypothesized that the GFP_11_ tag might be too far away from the VP30 binding site at 558-565 to complement GFP_1-9_ with VP30-GFP_10_. Consistent with this hypothesis, deleting amino acids 855-1720 yielded a construct that efficiently complemented GFP1-9 when co-expressed with VP30-GFP10. Further deletion of residues 502-626, which removed the VP30 binding site, reduced GFP complementation to background levels. Neither RBBP6-GFP11 construct complemented GFP1-9 fluorescence when co-expressed with VP40.

**Fig. 10.**
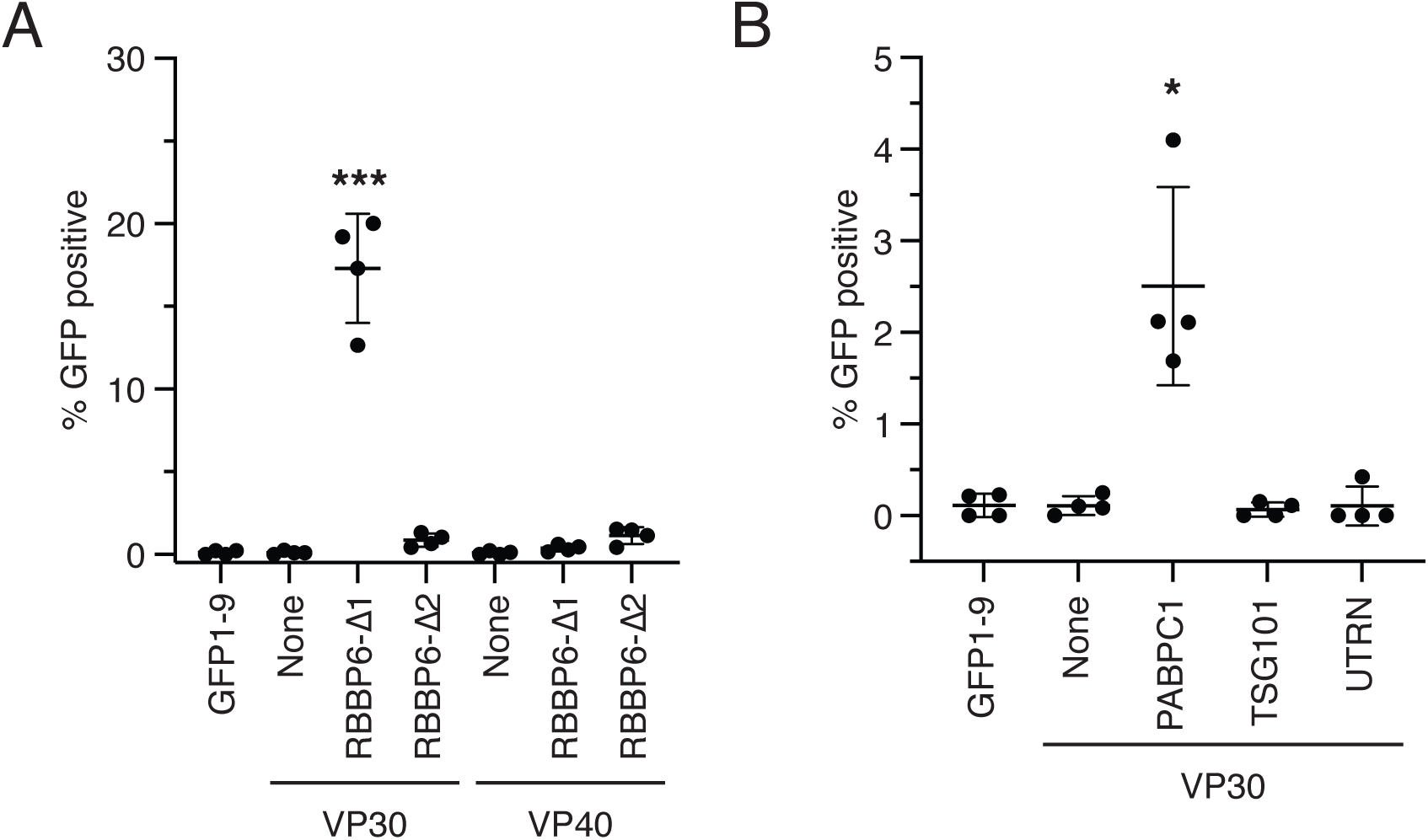
EBOV VP30 interacts with RBBP6 and PABPC1 in a tripartite split-GFP assay. A. HEK293 cells were transfected with plasmids expressing GFP_1-9_-T2A-mCherry, VP30-GFP_10_ or GFP_10_-VP40, and RBBP6-Δ1-GFP_11_ (deletion of amino acids 855-1720) or RBBP6-Δ2-GFP_11_ (deletions of amino acids 502-626 and 855-1720); None indicates no GFP_11_ plasmid was included. GFP cells were imaged using a PerkinElmer Opera Phenix High Content Screening System and quantified as a percentage of mCherry-positive cells. B. HEK293 cells were transfected with plasmids expressing GFP_1-9_-T2A-mCherry, VP30-GFP_10_, and PABPC1-GFP_11_, TSG101-GFP_11_, or UTRN-GFP_11_, and imaged as described for part A. * p value < 0.05, *** p value < 0.001, as determined by one way ANOVA with multiple comparisons.

Having established the split-GFP system for VP30, we co-expressed PABPC1 tagged at the C-terminus with GFP_11_ with GFP_1-9_ and VP30-GFP_10_ (Fig. 10B). As negative controls we co-expressed VP30-GFP_10_ two proteins, TSG101-GFP_11_ and UTRN-GFP_11_, that efficiently complemented VP40 in the tripartite GFP system but that are not known to bind to VP30 (40). PABPC1-GFP_11_, but not TSG101-GFP_11_ or UTRN-GFP_11_, caused a significant increase in fluorescence from GFP_1-9_ when co-expressed with VP30-GFP_10_, providing additional support for an interaction between VP30 and PABPC1.

### PABPC1 is recruited to inclusion bodies in EBOV-infected cells

Since VP30 binds to PABPC1 in uninfected cells, we next investigated whether EBOV infection altered the localization of PABPC1 (Fig. 11). HeLa cells were infected with EBOV expressing VP30 with a C-terminal FLAG epitope tag and fixed at 18 hpi, then stained for viral proteins and RNA. PABPC1 was evenly distributed throughout the cytoplasm of infected and uninfected cells. However, PABPC1 accumulated in intensely stained puncta separated by areas with little or no detectable staining within VP35-labelled inclusion bodies. Viral RNA co-localized within these regions of intense PABPC1 staining. VP30 was distributed throughout the interior of the inclusion body, including regions containing PABPC1.

**Fig. 11.**
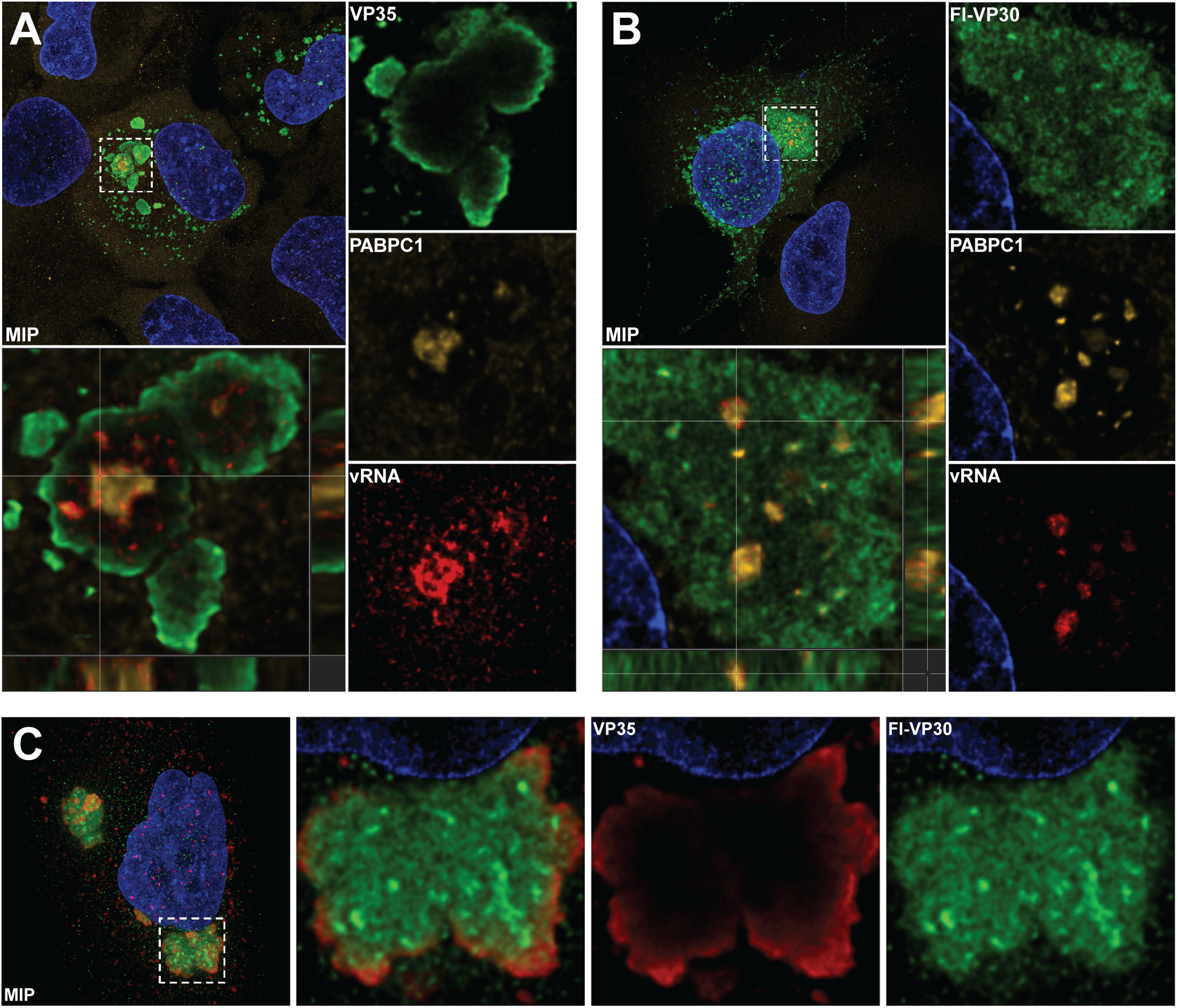
PABPC1 is concentrated in inclusion bodies in EBOV-infected cells. A) HeLa cells were challenged with rEBOV expressing VP30-FLAG at MOI 0.3 and fixed 18hpi. Cells were labeled with VP35 (green), PABPC1 (yellow), and NP vRNA (red). A max intensity projection (MIP) is shown on top left, and orthogonal view shown on bottom left. Single plane images of an inclusion body are shown on the right. B) As in A, EBOV infected cells were labeled with VP30 (green), PABPC1 (yellow), and NP viral RNA (red). A max intensity projection (MIP) is shown on top left, and orthogonal view shown on bottom left. Single plane images of an inclusion body are shown on the right. VP30, PABPC1, and NP vRNA signal overlaps in distinct puncta inside the inclusion bodies. C) To examine VP30-FLAG localization to inclusion bodies EBOV infected cells were labeled with VP35 (red) and VP30 (green) in EBOV infected cells 18hpi.

### PABPC1 is required for early EBOV replication

To determine if VP30 was required for EBOV replication, we inhibited expression of PABPC1 by RNA interference. We first evaluated four PABPC1 siRNAs for their ability to inhibit PABPC1 expression (Fig. S7). We then used two siRNAs to inhibit PABPC1 expression and assessed their effects on EBOV replication (Fig. 12). At 16h post infection, both PABPC1 siRNAs significantly reduced the number of infected cells as measured by RNAFISH with probes targeting NP and VP35 mRNA. To confirm this reduction, the level of NP mRNA was measured via RT-qPCR and was found to be significantly reduced in PABPC1 depleted cells, indicating that PABPC1 was required for EBOV infection.

**Fig. 12.**
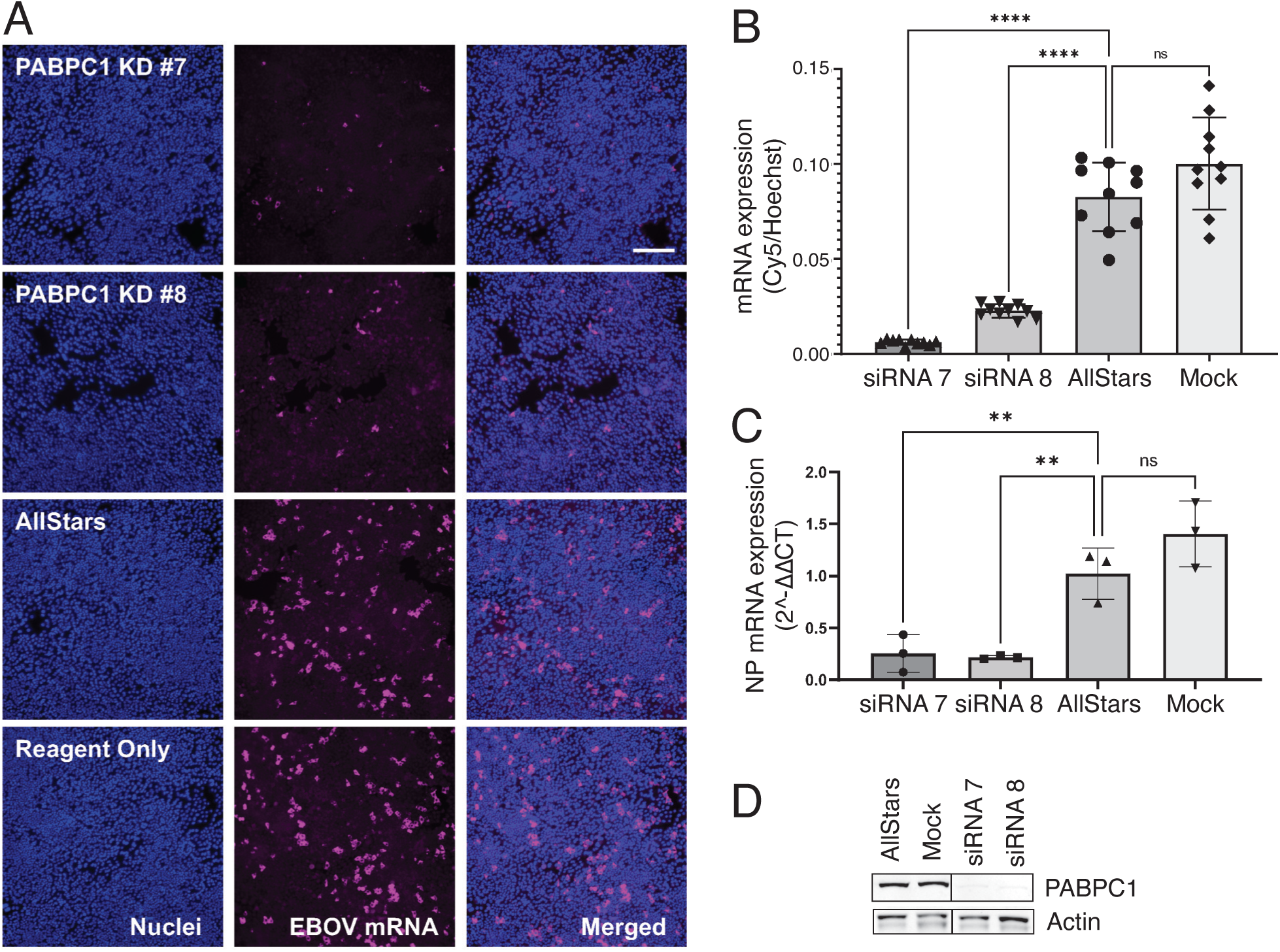
Inhibition of PABPC1 expression reduces EBOV replication. HeLa cells were mock-transfected (reagent only) or transfected with two distinct PABPC1 siRNAs or nontargeting AllStars negative control siRNAs. At 48 h post-transfection, cells were infected with EBOV at an MOI of 0.1. A.) At 16 hpi, samples were inactivated in 10% neutral-buffered formalin and RNAFISH was performed using probes detecting (+) sense NP and VP35 RNA (magenta). Nuclei were visualized by staining with Hoechst (blue). Scale bar = 250 μM. B.) Quantification of NP and VP35 RNA staining in panel A was performed in ImageJ by calculating the area occupied by mRNA signal and normalizing it to the area occupied by Hoechst (nuclei) signal. C.) In a parallel set of samples, total RNA was isolated by lysing the cells with Trizol reagent at 16 hours post infection. NP RNA levels were quantified by one-step RT-qPCR using a (+) sense NP probe to detect EBOV RNA and are normalized to the level of β-Actin RNA. D) Depletion of PABPC1 in siRNA-treated cells. A parallel set of samples were subjected to SDS-PAGE followed by western blotting with PABPC1 and actin antibodies. Molecular weight markers in kDa are shown at left.

## Discussion

This study expands the current understanding of EBOV–host protein interactions by identifying hundreds of previously unreported interactions using a large-scale Y2H approach. While no single method can capture the entire interactome, combining complementary strategies—such as Y2H, complex purification, and proximity-dependent biotinylation—provides a more comprehensive view of viral–host protein networks. Our findings overlap substantially with prior reports but also reveal novel interactions that may inform EBOV replication and pathogenesis.

To enhance PPI discovery, we employed libraries derived from infection-relevant tissues (liver) and cell types (macrophages), screened both full-length proteins and domain fragments, and retested over 200 interactions. Although the liver and macrophage libraries had more overlap than expected, the macrophage library yielded substantially more interactions and differentiates this work from other studies of EBOV-host protein interactions that typically use HEK293 or Huh7 cells. Use of multiple partially overlapping EBOV constructs to screen the library increased confidence in the interactions found in multiple independent screens and in many cases enabled the domain that mediated the interactions to be defined. Retesting a subset of interactions using a quantitative nanoluciferase reporter strain demonstrated that a high percentage of the interactions were reproducible in the Y2H assay. Finally, by incorporating data from our primary Y2H screens, Y2H retest assays, and representations in the cDNA libraries, we assigned confidence scores to all interactions from the study.

Use of Y2H AD libraries made of cDNA fragments allowed the human protein interaction domain to be delimited for some interactions. In the best case, overlapping clones from the Y2H screens precisely revealed the known binding site of VP30 on RBBP6. Surprisingly, this approach revealed a noncanonical VP30 binding site in NUFIP2. To date, all characterized VP30 binding sites are proline-rich and include a conserved tyrosine at the 3’ end (PPPPxY). However, this consensus sequence was absent from the VP30 binding site on NUFIP2 predicted by overlapping clones from the Y2H screens (346-DNSSAKIVPKISYASKVKENLNKTIQNSSVSP-377). Since mutations in VP30 that disrupted binding to NP and RBBP6 also abrogated interaction with NUFIP2, VP30 most likely binds to NUFIP2 through its NP/RBBP6 binding site in the structured C-terminal domain. VP30 interactions with other human proteins that lacked PPPxY consensus sites, including PABPC1 (see below), were also disrupted by mutations in the RBBP6/NP binding site. MEME analysis did not reveal alternative consensus sites in this set of VP30-binding partners, suggesting that VP30 has the potential to bind to a variety of non-proline-rich sequences, at least in the Y2H assay.

Integrating the EBOV-human protein interactions with EBOV-host cell functional interactions revealed human proteins that bind to EBOV proteins and that impacted EBOV replication or EBOV-induced c-Jun nuclear localization. In particular, we identified multiple interactions that link EBOV RNA stability and translation efficiency. We confirmed one of these interactions between VP30 and PABPC1 in multiple independent assays and showed that PABPC1 was concentrated in intensely stained puncta within inclusion bodies that overlapped with viral RNA. VP30 was found throughout the interior of inclusion bodies, including regions of PABPC1 staining, whereas VP35 was limited to the exterior. Inhibiting PABPC1 expression using siRNAs reduced VP35 RNA levels and virus replication. PABPC1 knockout in the EBOV OPS study reduced VP35 RNA levels in both the primary screen and a focused secondary screen (51). However, PABPC1 knockout showed different effects on VP35 protein levels in the primary and secondary screens, with no effect seen in the primary screen and proportional decrease in VP35 protein levels relative to VP35 RNA observed in the secondary screen (Fig. 3 and S8). The basis for this discrepancy is not clear but may be due to differences in timing (28 hpi in the initial screen versus 24 hpi in the secondary screen) or the sgRNAs used.

Despite being identified shortly after the discovery of poly(A) tails, the precise molecular functions of PABPC1 are still being uncovered (79). PABPC1 requires approximately 12 A residues for high affinity interaction, buts cover approximately 30 bases so most mRNAs will be bound to two copies of PABPC1 (80–82). Paradoxically, PABPC1 stabilizes mRNAs but also binds to and stimulates the activity of the enzymes that shorten poly(A) tails (deadenylases PAN1 and PAN2 and the CCR4-NOT complex) (83–86). In embryos, PABPC1 boosts translation by facilitating formation of a closed loop between the 3’ and 5’ ends through association with eIF4G at the 5’ cap (87–90). However, in differentiated cells, PABPC1 primarily affected mRNA abundance with no significant effect on the translational efficiency (91, 92).

Many viruses target PABPC1 for degradation to disrupt translation of cellular proteins (reviewed by (77)). Other viruses recruit PABPC1 to their mRNAs via interactions with viral proteins to promote translation (93, 94) or facilitate circularization of viral mRNAs by stabilizing interactions between PABPC1 and cap-binding translation initiation factors (95). EBOV’s interactions with PABPC1 are most similar to those of respiratory syncytial virus (RSV). PABPC1 binds to RSV nucleoprotein (N) and M2-1, which is functionally analogous to VP30 (96, 97). During RSV infection, PABPC1 accumulates in inclusion bodies and forms biomolecular condensates referred to as inclusion body associated granules (IBAGs) (98). IBAGs contain viral mRNA (but not viral genome) and eukaryotic initiation factors, which are recruited in an M2-1 and PABPC1-dependent manner (97–99). These observations led to a model in which IBAGs promote the assembly of ribonucleoprotein complexes with viral mRNAs to facilitate efficient translation of viral proteins (97–99). Morphologically, the PABPC1 puncta in EBOV inclusion bodies resemble those in RSV, providing the first evidence that EBOV also forms IBAGs. Whether these PABPC1 structures play a similar role in EBOV-infected cells remains to be determined.

In addition to binding poly(A) RNA, PABPC1 complexes with many cellular proteins, including NUFIP2 (100, 101) and CSDE1 (also known as UNR) (102, 103). CSDE1 can positively or negatively regulate cap-independent translation of cellular and viral RNAs through interactions with internal ribosome entry site (IRES) elements and multiple cellular proteins (104–108). CSDE1 also regulates mRNA stability in an IRES-independent manner in conjunction with its primary binding partner, STRAP, through translation-coupled mRNA turnover (109–111). Translating ribosomes displace mRNA-bound CSDE1 and STRAP, which stabilizes the mRNA and increases protein synthesis (109, 111). Loss of either CSDE1 or STRAP increased BACH1 mRNA levels but did not increase Bach1 protein to a similar extent (110). A very similar phenotype was observed in both CSDE1 and STRAP knockout cells in the EBOV OPS study, with a significant increase in VP35 RNA levels but little or no increase in VP35 protein in both primary and secondary screens (Fig. 3 and S8) (51). Together, these observations suggest that VP35 RNA levels are regulated by CSDE1 and STRAP, likely by coupling to translation in a manner similar to that of c-Fos and Bach1 mRNAs. EBOV may further manipulate this regulatory mechanism through direct interactions with viral proteins. Our Y2H screen identified a high confidence interaction of CSDE1 with VP24 and a low confidence interaction with VP35. Carlson et al found that STRAP is proximal to VP35 protein in infected cells but did not demonstrate a direct interaction (51).

Knockout of PDCD2 had the opposite effect of CSDE1 and STRAP in the EBOV OPS study (51). In PDCD2 knockout cells, VP35 RNA levels were significantly reduced, but VP35 protein levels did not decrease proportionally. This suggests that either the remaining RNA was translated more efficiently in PDCD2 knockout cells or that only a portion the VP35 RNA in wildtype cells is capable of being translated and this population is somehow maintained in PDCD2 knockout cells despite the lower levels of RNA. PDCD2 is a ribosomal chaperone protein, which facilitates the formation of 40S ribosomes (112). PDCD2 binds to the 40S ribosomal protein RPS2 (uS5) co-translationally and traffics with it to the nucleolus but does not associate with mature 40S ribosomes (113, 114). Loss of PDCD2 expression increased the amount of free 60S ribosomes while decreasing the amount of 80S monosomes, an indication of a defect in 40S assembly (113, 114). This deficiency was partially corrected by overexpression of RPS2, suggesting PDC2 primarily functions through Rps2 (113). However, in the EBOV OPS screen, RPS2 knockout proportionally decreased both VP35 RNA and protein (51). PCDC2 knockout was more similar to that of another 40S ribosome chaperone, AK6, which dramatically reduced VP35 RNA levels but had little effect on VP35 protein (Fig. 3 and S8) (51, 115).

PDCD2 is one of five proteins (ANKMY2, EGLN1, PDCD2, SMYD2, and SMYD3) that contain a MYND domain and that bind to EBOV NP. Both PDCD2 and EGLN1 have been identified in multiple independent reports, strongly suggesting they are valid EBOV NP binding partners (21, 25, 28). Of the MYND domain proteins, only PDCD2 and EGLN1 were required for EBOV replication, though their phenotypes were different (51). EGLN1 knockout caused a proportional decrease in both VP35 RNA and protein, whereas loss of PDCD2 preferentially affected VP35 RNA levels, as discussed above. A putative MYND domain binding site is present in the NP IDR (Fig. 5), which was required for binding to SMYD3 (24). Together, these observations suggest that EBOV NP recruits MYND domain proteins to enhance replication.

Integrating the Y2H hits with c-Jun nuclear localization revealed that most knockouts that reduced nuclear Jun localization had a proportional effect on VP35 protein levels, suggesting that reduced Jun trafficking was due to lower EBOV replication (Fig. 3B). However, knockouts of eight genes identified in the Y2H screen significantly increased Jun nuclear localization with little or no effect on VP35 protein production. In particular, knockout of the VP24 partner OTUD5 (also known as Deubiquitinating Enzyme A or DUBA), caused one of the largest increases in c-Jun nuclear localization without impacting EBOV replication (116, 117). OTUD5 is a deubiquitinating enzyme that regulates both innate and adaptive immune responses (118–122), and may represent another target by which VP24 regulates the host immune response.

### Limitations and Future Directions

While Y2H provides valuable insights into direct protein-protein interactions, it does not capture interactions that depend on complexes of viral proteins or on post-translational modifications that occur only in infected cells. In some cases, a single cellular protein interacted with multiple viral proteins, which could impact the effects gene knockouts on virus replication. Similarly, inhibiting or disrupting expression of cellular genes may influence virus replication indirectly by altering the expression of other cellular genes. Further validation in infected cells and mutational analyses to disrupt specific interactions, such as VP30-PABPC1 or VP24-CSDE1, will be essential to confirm functional relevance and to determine whether these pathways can be targeted pharmacologically.

## Acknowledgements

Funding support was provided by NIH/NIAID grants R01AI114814 and P01AI120943 to D.J.L. and R.A.D.

---

Fig. S1. EBOV bait constructs used for Y2H cDNA library screens. Each EBOV full length gene and the indicated gene fragments were fused to the 3’ or 5’ end of the Gal4 DNA binding domain (BD). Numbers below each EBOV protein indicate amino acid positions. Bait number refers to our internal nomenclature used to track each construct. First and third numbers (when present) indicate constructs fused to the C-terminus of the Gal4 DNA binding domain (BD), whereas second and fourth numbers indicate fusions to the N-terminus of the BD. All constructs were cloned into pOBD7A, except 1-16, which were cloned into pOBD2 or pOBD4 for 3’ and 5’ fusions to the Gal4 AD. Further details can be found in Table S1.

Fig. S2. Retest of EBOV-human gene pairs in BK100Nano. Graphs show the mean relative nanoluciferase activity normalized to the OD600 of the culture for pairs of EBOV and human proteins identified in Y2H cDNA library screens (black dots) and the normalized relative luciferase activity from individual replicates of each EBOV bait construct with AD-GFP (green dots). Red bar indicates the value of the mean relative nanoluciferase activity plus five standard deviations for the EBOV bait construct with AD-GFP.

Fig. S3. EBOV-human protein interaction network with confidence scores. Network of all EBOV-human protein-protein interactions identified in the EBOV Y2H screens. Interaction confidence is indicated by the color of the lines connecting the EBOV proteins (diamonds) to human proteins (ovals). Blue lines, high confidence interactions (group 1); Black lines, medium confidence (group 2); low confidence, gray lines (group 3).

Fig. S4. Overlap with published EBOV-human protein-protein and functional interaction studies. Subnetwork of EBOV-human protein-protein interactions identified in published studies (EBOV proteins, diamonds; human proteins, ovals). Interactions identified in previous studies are indicated by blue lines. Human genes identified in functional studies are indicated by light blue ovals. Additional details can be found in Tables S5-S7.

Fig. S5. EBOV-human protein interactions mapped to viral protein domains. Human proteins that interacted with EBOV proteins are shown associated with the smallest protein domain that identified the interaction. Human proteins that were identified by other larger domains are shaded as indicated in the figure.

Fig. S6. Poly-proline motifs in human proteins that bind to EBOV VP30. Human proteins are shown as gray bars whose sizes are proportional to the number of amino acids, except for HERC, which has been reduced 50%. Poly-proline sequences identified by MEME are shown as red bars. Height of the red bars indicates how well the sequence matched the consensus motif identified by MEME.

Fig. S7. Effect of mutations in VP30 C-terminal domain on binding to human proteins. The same from Fig. 7 is plotted, except that the nanoluciferase values have not been normalized to wildtype VP30 for each human gene. Dot size corresponds to relative nanoluciferase activity. Wildtype VP30 constructs that interacted with each human gene are indicated in the first column (+, VP30 C-terminal domain fused that the N-terminus of BD; ++, VP30 C-terminal domain and VP30 full length fused that the N-terminus of BD; +++, VP30 C-terminal domain, VP30 full length fused that the N-terminus of BD, and VP30 full length fused that the C-terminus of BD.

Fig. S8. Inhibition of PABPC1 expression by RNA interference. HeLa cells were transfected with four distinct PABPC1 siRNAs or nontargeting AllStars negative control siRNAs. Lysates were prepared at 48 h post transfection and subjected to SDS-PAGE followed by western blotting with PABPC1 and actin antibodies. Molecular weight markers in kDa are shown at left.

Fig. S8. Integration of host factors from the EBOV Y2H screen and the secondary CRISPR-Cas9 knockout optical pooled screen. Late HeLa VP35 Protein Median Cell Mean cumulative delta AUC from the Carlson et al. secondary screen, Table S5, was plotted as a function of Late HeLa VP35 RNA FISH Median Cell Mean cumulative delta AUC (51). Gene knockouts that had a significant effect on both VP35 RNA and protein levels (p<0.0001) are shown in cyan. Select hits from the Y2H screen are labelled. Simple linear regression of all gene knockouts in the secondary screen showed a strong correlation between VP35 RNA and protein levels (R^2^ = 0.91, black line). Brackets show the difference between the observed VP35 protein levels and those expected based on RNA levels.

